# Cardiac Calsequestrin is a Physiological Dimer that Polymerizes through a Ca²⁺-Triggered Cooperative Switch

**DOI:** 10.64898/2026.04.04.716468

**Authors:** Chiara Marabelli, Demetrio J. Santiago, Emanuele Pirana, Chiara Di Antonio, Anselmo Canciani, Martino Bolognesi, Federico Forneris, Silvia G. Priori

**Affiliations:** Molecular Cardiology Dept., IRCCS Maugeri of Pavia, 27110 Pavia, Italy; Centro Nacional de Investigaciones Cardiovasculares Carlos III (CNIC), 28029 Madrid, Spain; Biology and Biotechnology Dept., University of Pavia, 27100 Pavia, Italy; Molecular Medicine Dept., University of Pavia, 27100 Pavia, Italy; Biosciences Dept., University of Milan, 20133 Milan, Italy

**Keywords:** calsequestrin, protein polymerization, electrostatic regulation, cardiac arrhythmia, catecholaminergic polymorphic ventricular tachycardia

## Abstract

Cardiac Calsequestrin (CASQ2) polymerizes within the junctional sarcoplasmic reticulum to buffer Ca²⁺ and regulate ryanodine receptor 2 (RyR2) gating, yet the molecular mechanism governing this process remains poorly understood. Using an integrated set of complementary approaches spanning single-particle biophysics, bulk solution measurements, and polymer chemistry, we demonstrate that CASQ2 is an intrinsic dimer at nanomolar concentrations and under physiological ionic conditions, independently of Ca²⁺. In addition, Ca²⁺-dependent polymerization operates as a highly cooperative switch between a stable oligomeric phase and a high-order polymeric state. Physiological amounts of K⁺ ions modulate this switch through a biphasic electrostatic mechanism, supporting polymerization at low concentrations and inhibiting it beyond charge neutralization (∼194 mM). These findings redefine CASQ2 as an intrinsic dimer with polymerization-switch properties, and provide a mechanistic framework for understanding how catecholaminergic polymorphic ventricular tachycardia type 2 mutations, distributed evenly across the CASQ2 surface, cause disease through two distinct pathological trajectories.

## Introduction

Calsequestrin (CASQ) is critical to the Ca^2+^-mediated process of excitation-contraction coupling (ECC) in skeletal and cardiac muscle cells. It is the principal Ca^2+^-binding protein in the lumen of the junctional Sarcoplasmic Reticulum (jSR), where it regulates either directly or indirectly the opening and refractoriness of the ryanodine receptor (RyR) Ca²⁺ channel, and thereby controls contraction initiation and termination. In vivo, CASQ function is intimately linked to its Ca^2+^-dependent assembly into filamentous electron-dense structures within the jSR^1–3^, that disassemble upon extreme Ca^2+^ depletion^4^. Loss of these luminal jSR polymeric structures is a characteristic of pathological conditions ^1,3,5^.

CASQ Ca^2+^-driven polymerization pairs with its low-millimolar affinity, high-capacity Ca^2+^-binding (up to 60 or 80 Ca^2+^ ions per molecule, depending on the isoform)^6^. Unlike typical Ca^2+^-binding proteins, CASQ lacks any EF-hand motifs or defined acidic Ca^2+^-binding site. At least one fourth of CASQ residues are glutamates or aspartates, whose charge neutralization by Ca^2+^ ions allows the compaction of its three thioredoxin-fold domains (Figure 1A). This globular core features a highly negative surface, and two flexible, structurally essential, terminal regions (Figure 1). The swapping of the N-terminal tails locks two CASQ monomers in the crystallographic dimeric (Figure 1B)^3,7–9^. In the traditional model of polymerization, Ca^2+^-mediated bridges allow both this “front-to-front” dimer, as well as additional “back-to-back” inter-dimer contacts proposed to drive polymer assembly. The highly acidic C-terminal tail (Figure 1C), shapes the isoform-specific kinetics of Ca²⁺-dependent polymerization and Ca²⁺-binding capacities^10–13^, yet its structural characterization, critical for understanding CASQ2/Ca^2+^ polymerization mechanism, is challenged by its high flexibility^12^.

**Figure 1.**
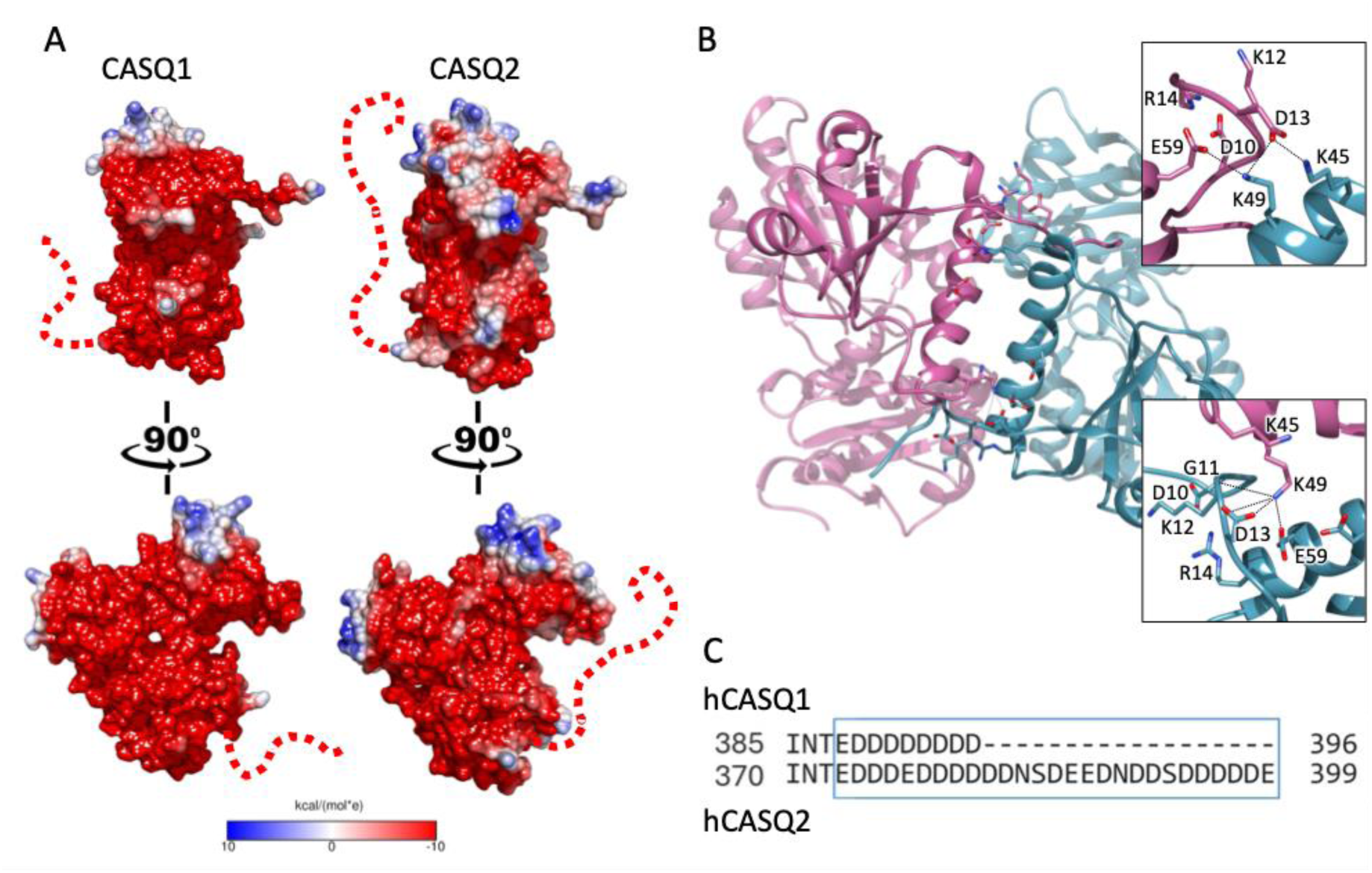
Structural and electrostatic comparison between CASQ1 and CASQ2. **A**) Electrostatic surface potential maps (±10 kcal/mol·e) of CASQ1 and CASQ2 show the highly negative charge distribution across both proteins. The intrinsically disordered C-terminal tail (dashed red line) includes extensive clusters of negative charges. B) Ribbon representation of the CASQ2 dimer model (magenta and cyan monomers), with insets showing zoomed views of the crystallographic interface regions shaped by the swapped N-terminal segments. **C)** Sequence alignment of the C-terminal regions of human CASQ1 and CASQ2 underscores the more extensive aspartate/glutamate enrichment in CASQ2.

CASQ polymerization is classically viewed as a Ca^2+^-driven sequential process, where monomer compaction enables dimerization, followed by tetramerization and the formation of a ribbon-like, linear polymers. However, this model does not justify the mechanism of the severe CASQ-related pathologies such as Tubular Aggregate Myopathies or Malignant Hyperthermia for the skeletal CASQ1 ^14–17^, or the highly lethal Catecholaminergic Polymorphic Ventricular Tachycardia type 2 (CPVT2) for the cardiac CASQ2 ^9,18–21^. Would specific interfaces be involved in the polymerization process, then pathological missense mutations would cluster at defined patches rather than being spread all over the protein surface.

The prevailing view on CASQ polymerization as a linear cascade of Ca^2+^-dependent events^1,8,12,22,23^ has been suggested mainly by *in vitro* turbidity measurements, which measure protein aggregation via light-scattering at 350 or 600 nm wavelengths, where absorption by proteins is minimal. Although turbidimetry captures the kinetics of protein multimerization, it cannot discriminate the nature of the light-scattering bodies, such as soluble aggregates or linear fibers^24,25^.

Also, while the properties of CASQ have been widely studied with reference to the effects induced by Ca^2+^ ^7–9,11,12,17,23,26^, the primary ion of interest in the jSR, the roles of other relevant cations, such as K^+^ and Mg^2+^ have been far less evaluated. At the same time, recent findings reveal that low millimolar Mg^2+^ can unmask differences in Ca^2+^-responsiveness across distinct CASQ2 variants^9,23^. Hence, the potential exists to incorporate the role played by other physiologically relevant ions in modulating CASQ folding and polymerization.

In this broad context we comprehensively examined the interplay between ionic strength, protein concentration, and Ca^2+^ abundance in shaping the polymerization of the cardiac isoform CASQ2. Using a combination of approaches from single-molecule biophysics and polymer chemistry, we show for the first time that dimerization is an intrinsic property of cardiac CASQ2 rather than being exclusively Ca²⁺-dependent. Moreover, our data identify a previously uncharacterized Ca²⁺-dependent switch-like transition from oligomeric to polymeric forms, challenging the traditional view of CASQ polymerization as a stepwise process. Our data indicate that CASQ2 dimers may identify the physiologically relevant, functionally active unit, which responsiveness to Ca^2+^ integrates additional factors beyond Ca²⁺ abundance.

## Results

### Physiological quantities of K^+^ ions support CASQ2 conformational compaction

Published NMR and CD spectra ^10,16,27–31^ show that CASQ2 secondary structure is extremely sensitive to the ionic conditions, and that Ca^2+^ is orders of magnitude more efficient than monovalent cations in promoting CASQ2 molecular compaction. Under low ionic strength conditions (e.g. 20 mM NaCl), 0.3 mM Ca^2+^ largely stabilizes CASQ secondary structures content, an effect that is matched by three orders of magnitude higher quantities of K^+^ or Na^+^ (300 mM). The mechanistic interpretation is that CASQ retains a molten globule state until its negative charges are adequately masked by levels of free Ca^2+^ comparable to those found within the jSR of muscle cells (0.3-1 mM) ^32,33^. However, K^+^ is usually present within 120-225 mM range in the cardiomyocyte cytoplasm, whereas the cytosolic concentration of the second most abundant monovalent cation, Na^+^, falls in the 5 to 15 mM range^32^. In light of this, and of the accumulating evidence on the ionic-sensitive nature of CASQ2 Ca^2+^-dependent polymerization as well^9,23^, we sought to investigate the role of the physiologically abundant jSR ions on CASQ2 conformational equilibrium. This study focuses on the effect of K^+^ ions on CASQ2 Ca^2+^-dependent polymerization kinetics, on the generally observed assumption that CASQ2 is exposed to higher amounts of K^+^ rather than Na^+^ ions.

To dissect how K^+^ ions affect the conformational compaction of CASQ2 molten globule state, we measured the apparent volume of the CASQ2 monomer by mass photometry (MP), in the presence of varying KCl concentrations. The MP method allows to calculate the size of CASQ2 indirectly, from the light scattering variation produced by single particles in solution that stochastically encounter a measurement surface. As for other experiments, the absence of contamination from Ca^2+^ or other divalent ions was ensured by dialysis in EDTA prior to measurements. Our MP data confirm the expected proportional collapse of CASQ2 size with respect to the ionic strength (Figure 2A, Figure S1). Increasing KCl levels, drive an evident compaction of CASQ2 monomer, maximal in the 150-200 mM range,where the difference with apparent solution volume observed in 25 mM KCl is 6.4 ± 2.4 kDa (Figure 2A). Supporting the cited K^+^/Ca^2+^ cooperativity and higher efficacy of Ca^2+^ in CASQ2 folding, a similar effect is also triggered by 10 mM CaCl_2_ in 50 mM KCl (Figure S2). Notably, the addition of millimolar amounts of CaCl_2_ in 200 mM KCl does not trigger any further compaction (Figure S2). The inverse relationship between ionic strength and CASQ2 hydrodynamic size is rather reversed beyond 200 mM (Figure 2A), suggesting that extensive charge screening may perturb either CASQ2 conformational architecture and/or its hydration shell.

**Figure 2.**
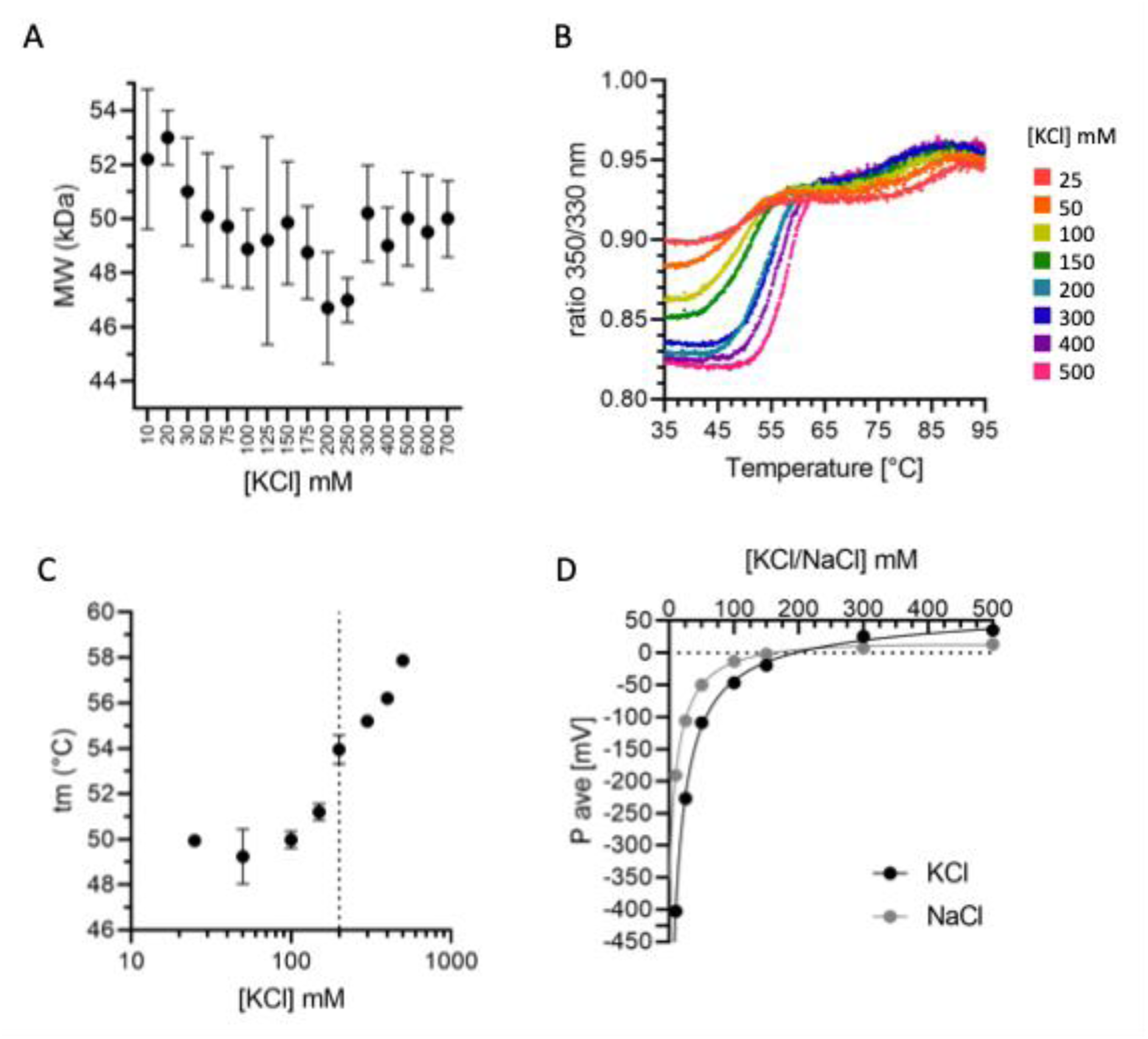
Monovalent cations modulate the tertiary structure and solvation shell of CASQ2. **A)** CASQ2 size distribution at varying KCl concentrations as determined by MP. Higher ionic strength promotes the compaction of the monomer observed at 200 mM KCl (dotted line). Data are presented as median + range from at least three independent measurements. **B)** Thermal denaturation curves for CASQ2 samples under varying KCl concentrations, from which t_m_ values are extrapolated. Data are presented as mean from three independent measurements. **C)** The value of CASQ2 t_m_ increases rapidly with KCl concentrations up to 200 mM (dotted line). At concentrations higher than 200 mM KCl, the t_m_ increases further. Data are presented as mean ± SD from three independent measurements. **D)** ζ-potential measurements of CASQ2 at increasing KCl (black circles) and NaCl (grey circles) concentrations show a progressive reduction in surface charge. At 10 mM KCl and NaCl, the surface potential of CASQ2 is on average respectively -403 mV and -191 mV. Both KCl and NaCl neutralize the protein at similar concentrations. At 500 mM KCl and NaCl, the ζ-potential of CASQ2 is respectively on average +35 mV and +13 mV. Data are presented as mean ± SD from three independent measurements.

We next explored thermal denaturation assays as a specular method to test the ionic dependence of CASQ2 conformation. By tracking the change in the intrinsic fluorescence of the protein’s tryptophan residues (usually buried in a solvent-inaccessible environment) at increasing temperatures, thermal denaturation assays measure the kinetics of their exposure to the aqueous solvent, usually mirroring the kinetics of thermally-induced protein unfolding. The temperature at which the highest slope of the denaturation curve is registered defines the melting temperature (t_m_). Our analyses of CASQ2 thermal stability in varying KCl concentrations (Figure 2B) reveal a trend similar to that observed for size compaction: with increasing KCl levels (from 25 to 200 mM), the increase in t_m_ and the variation in fluorescence emission spectra are indicative of a tightening of the hydrophobic core. An inflection in these trends is registered at 200 mM KCl, a condition in which the t_m_ of CASQ2 is higher by 4 °C than that registered in 50 mM KCl (Figure 2C). Taken together, both the volume variation measured by MP (Figures 2A, S2), and the thermal stabilization (Figures 2B-C) demonstrate that 200 mM K^+^ concentration maximally supports the compaction of CASQ2 tertiary structure.

Counterintuitively, both the apparent volume occupied by CASQ2 (Figures 2A, S2) and its hydrophobic core compaction (Figures 2B-C) rise when the ionic strength increases above 200 mM KCl (Figures 2A, 2C). To interpret these apparently contrasting results, we investigated CASQ2’s zeta (ζ)-potential, a proxy for exposed surface charges and ionic solvation shell of a particle in solution (Figure 2D). In 10 mM KCl, the abundant CASQ2 negative charges account for a highly negative ζ-potential of -400 mV, and become fully neutralized (0 mV) at the physiological concentration of 194 mM [K⁺] ^32^. Functionally, this implies that in the cellular context, even in the absence of Ca^2+^ ions, CASQ2 negative charges are mostly or completely shielded, allowing proper hydrophobic collapse and conformational packing of the CASQ2 polypeptide chain. The ζ-potential versus [K⁺] curve is well fit (R² = 0.9990) by a one-site saturation model, hence a site-specific neutralization mechanism where K⁺ ions bind directly to a defined number of locations on the acidic CASQ2 surface, rather than diffuse screening. A similar saturable behavior is observed with Na^+^ (R² = 0.9996, Figure 2D), though Na⁺ is nearly twice more effective than K⁺ at neutralizing the protein’s negative surface charge at lower than 100 mM concentrations due to its higher charge density. At higher equimolar levels, both ions ultimately cause charge reversal, (positive ζ-potential values of +35 mV and +14 mV at 500 mM KCl and NaCl, respectively), a typical consequence of ion accumulation over densely charge surfaces beyond isoelectric neutralization, which also pairs with the formation of a thicker solvation shell in solution^34^. This effect, observed for KCl concentrations above 200 mM, is consistent with the parallel increase in the apparent molecular volume of CASQ2 monomers (Figure 2A) and with the reduced solvent penetration into the hydrophobic core (Figures 2B-C). Together, all measurements on CASQ2 ion-dependent conformational fluctuations collectively indicate the 194-200 mM [K⁺] condition as a critical switch point between insufficient neutralization the abundant negative charges (which accompanies tertiary folding), and their excessive screening.

### CASQ2 forms dimers at physiological KCl concentrations in the absence of Ca^2+^

From ζ-potential curves it appears that CASQ2 is at least partially neutralized at the physiological concentrations of K^+^ of 140-225 mM ^32^. Masking of the abundant inter-repulsive negative charges not only facilitates polypeptide folding, but would also allow inter-monomer interactions, even in absence of Ca^2+^. Notably, most of the crystallographic dimeric structures have been solved in the absence of any divalent cation^3,14^. We next explored whether K⁺ ions alone (i.e. in the absence of Ca^2+^) can support quaternary assembly of CASQ2. Size-Exclusion Chromatography (SEC) profiles of CASQ2 under varying ionic conditions (Figures 3A, S4), indeed reveals the existence of multiple forms of Ca^2+^-free CASQ2 oligomers (dimers, trimers, and tetramers) with distinct sensitivities to the ionic strength: the dimer persists across all conditions, whereas the previously unreported trimeric species vanish beyond 50 mM KCl, and tetrameric ones peak at 100 mM KCl. Batch-SAXS analysis of CASQ2 revealed an identical biphasic dependency on the ionic strength. At concentrations similar or lower than those tested in SEC ([CASQ2] ≤ 56 𝜇M, Figure 3B, S5), the radius of gyration (R_g_) of the assemblies increases and falls with a peak at 100 mM KCl. This trend, and the analysis of the Guinier region (Figure S6), greatly support the electrostatic nature of the CASQ2 quaternary assemblies: (i) at 50 mM KCl the repulsion between highly negative surfaces is still too strong to allow inter-particle interaction, (ii) from here, increasing [KCl] initially favor sticking between oppositely charged patches, (iii) yet when approximating charge neutralization, also these sites are masked, and the thicker solvation shell increases the entropic cost of interaction.

**Figure 3.**
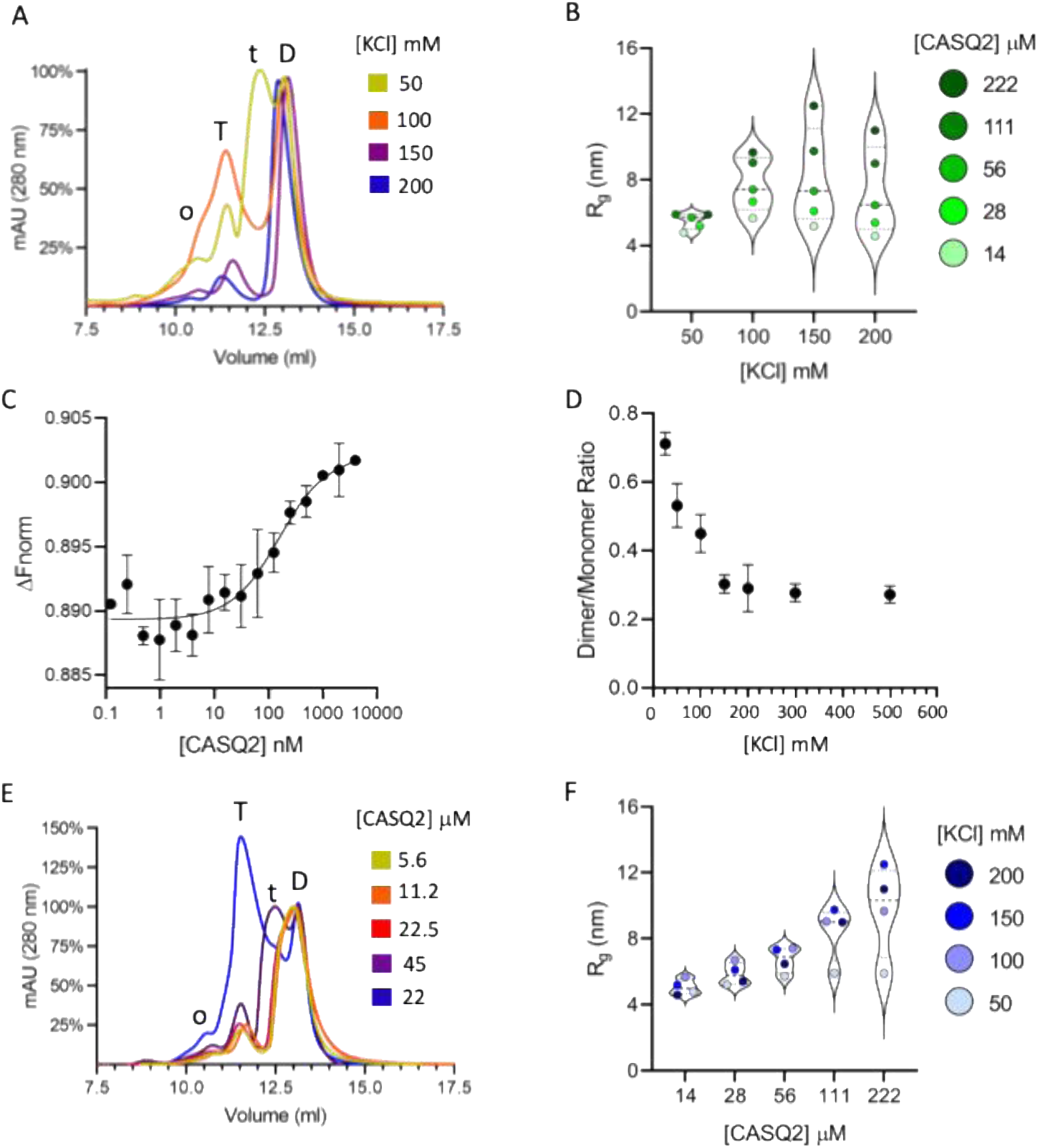
CASQ2 features an oligomerization propensity which is independent of the presence of Ca^2+^. **A)** SEC analysis of CASQ2 (45 µM) (Superdex 200 10/300 GL column) at increasing KCl concentrations (50 mM, 100 mM, 150 mM, and 200 mM). The identity of elution peaks to dimers (D), trimers (t), tetramers (T), and oligomers (o), is based on the expected molecular weight of the eluted species, calculated from the column size calibration curve (Figure S5). **B)** SAXS analysis of CASQ2 particles Rg in solution Vs. [KCl], with datapoints grouped by [KCl]. Coloring (light-dark green) highlights increasing [CASQ] within the subsets. Dashed lines show medians, dotted lines show interquartile ranges. **C)** MicroScale Thermophoresis (MST) analysis of the concentration-dependent self-association of CASQ2 in PBS buffer (containing 137 mM NaCl and 2.7 mM KCl). The increasing thermophoretic signal indicates that CASQ2 begins to form higher-order assemblies already in the nanomolar range. Data are presented as mean ± SD from three independent measurements. **D)** Quantification of dimer-to-monomer ratio from mass photometry (MP) across increasing [KCl], showing that a portion of CASQ2 dimers is significantly destabilized at higher than 100 mM salt concentrations. **E)** SEC profiles of CASQ2 samples (Superdex 200 10/300 GL column, see Figure S5) at varying protein concentrations (5.6 to 225 µM) running in a Ca^2+^-free, 50 mM KCl buffer, showing progressive shift from monomers to dimers (D) and tetramers (T) in a concentration-dependent manner. The oligomeric species (o) eluted at volumes accounting for an apparent mass of 200 ± 17 kDa, consistent with CASQ2 pentamers. **F)** SAXS analysis of CASQ2 particles Rg in solution ionic strength VS [CASQ2], showing single datapoints grouped by protein concentration. Data color (light-dark blue) highlights increasing [KCl] within the separate [CASQ2] subsets. Dashed lines show medians, dotted lines show interquartile ranges.

A second relevant observation from SAXS experiments, which well-aligns with SEC data, is that the species assignable to a dimer (R_g_= 4-6 nm, mainly detected at 14 𝜇M CASQ2 and/or at 50 mM KCl) is consistently present across ionic conditions, and even shrinks with increasing [KCl]: a behavior mirroring the ion-dependent conformational packing of the protein (Figure 2). The existence of ionic-independent dimeric species among multiple electrostatic assemblies, prompted us to analyze further the CASQ2 self-assembly at physiological ionic concentration (PBS buffer: 137 mM NaCl and 2.7 mM KCl) by MicroScale Thermophoresis (MST). MST quantifies how the thermal-dependent increase in the mobility of fluorescently labeled molecules in solution changes in the presence of a binding partner. A constant amount of fluorescently labeled CASQ2 was incubated with varying concentrations of non-labeled CASQ2. The first observation brought by MST experiments is that, in a physiological ionic condition and in absence of its canonical ligand, Ca^2+^, CASQ2 can self-assemble already at nanomolar levels (Figure 3C). MP experiments further confirmed this insight: CASQ2 forms KCl- sensitive, Ca^2+^-free dimers, which proportion with is diminished by KCl increasing between 50 and 150 mM K^+^ levels, and remains stable for higher ionic strength conditions (Figures 3D, S1, S3). This is consistent with the interpretation that a fraction of Ca^2+^-free CASQ2 dimers is resistant to ionic strength, whereas trimers and tetramers are lower-affinity electrostatic assemblies that dissolve with increasing surface charge screening.

However, what also appears evident from MST is that CASQ2 self-assembly does not plateau at a saturating concentration, but instead increases steadily, extending into the high micromolar regime (Figure S7). Such continuous signal shift once more suggests that CASQ2 can form supra-dimeric assemblies via multiple weak, non-exclusive interfaces, where the probability of forming productive interactions almost linearly depends on the number of available sites (solvent exposed charges and protein concentration). This is consistent with both SEC (Figure 3E), SAXS (Figure 3F), and MP (Figure S3B-C) data, which show a progressive shift of the quaternary species toward larger assemblies (lower elution volumes or larger R_g_ / apparent volume), that is proportional to the experimental concentration. Of note, no high-order polymer was observed by any of these methods. Turbidimetric inspection of 2.5 μM CASQ2 under varying ionic conditions did not reveal the presence of any large, light-scattering species (Figure S8). Taken together, these findings indicate that CASQ2 undergoes Ca²⁺-free self-association under physiological ionic conditions through a continuum of weak, multivalent interactions. Among the multiple electrostatic assemblies observed, a Ca^2+^-free dimeric assembly form emerges as the most stable species across ionic strength conditions.

### The CASQ2 Ca^2+^-free oligomers compete with Ca^2+^-driven quaternary assemblies

The co-existence of ionic-insensitive Ca^2+^-free CASQ2 dimers alongside electrostatic forms of assembly, raised the question on their specific contribution to the Ca^2+^-driven polymer nucleation. To this aim, we tested via MP the effect of Ca^2+^ on 10 nM CASQ2 and in 50 mM KCl (i.e. a condition in which the dimer is not dominant with respect to the monomer, Figure 4A). In line with a purely electrostatic effect, increasing amounts of Ca^2+^ shifted the equilibrium toward formation of dimers (Figure 4A), which become particularly evident at 20 mM and higher Ca^2+^ concentrations. However, and in disagreement with a direct consequence of lowering the inter-monomer repulsive forces, this growth is not immediate: for intermediate Ca^2+^ levels (1-10 mM), there is a subtle dip in the dimer-to-monomer ratio (Figure 3A). This effect becomes even more prominent at 25 nM CASQ2 (which favours CASQ2 electrostatic assembly, Figure 4B), suggesting a competition between electrostatic forms of the dimer and Ca^2+^-induced ones. By inhibiting electrostatic forms of dimerization in 200 mM KCl, the formation of Ca^2+^-dependent dimers is not inhibited anymore, but rather supported in the 1-10 mM Ca^2+^ range (Figure 4B). These data suggest that Ca^2+^ may promote quaternary interactions even when the protein charges are completely neutralized, and that Ca^2+^-induced CASQ2 dimeric states are in competition with electrostatic interactions (blue points in Figures 4B). We next used SEC to assess the Ca^2+^-responsivity of CASQ2 Ca^2+^-free species formed at higher concentrations under increasing ionic conditions (Figure 4C-F). Of note, at 45 μM CASQ2, no monomeric species is detectable (as in Figures 3A-B), and the net effect of Ca^2+^ is rather an increase of the tetrameric Vs. the dimeric species. Mirroring what observed in MP, the effect of Ca^2+^ (here tetramerization) is more prominent when electrostatic assemblies (i.e. tetramers and trimers in 50 and 100 mM KCl, Figures 4C-D) are maximally inhibited at 200 mM KCl (Figure 4F). Taken together, MP and SEC analyses clearly indicate that the Ca^2+^-driven quaternary assembly of CASQ2 does not build upon pre-existing electrostatic assemblies, but is rather in competition with them. Taking into account the physiological concentration of the CASQ2 protein within the jSR compartment (estimated to fall in the 1-4 mM range, corresponding to 50-200 mg/ml)^3,6,22^, the ionic environment and its variations would be highly relevant in shaping the equilibrium between the highly electrostatic nature of CASQ2 and its Ca^2+^-dependent polymerization.

**Figure 4.**
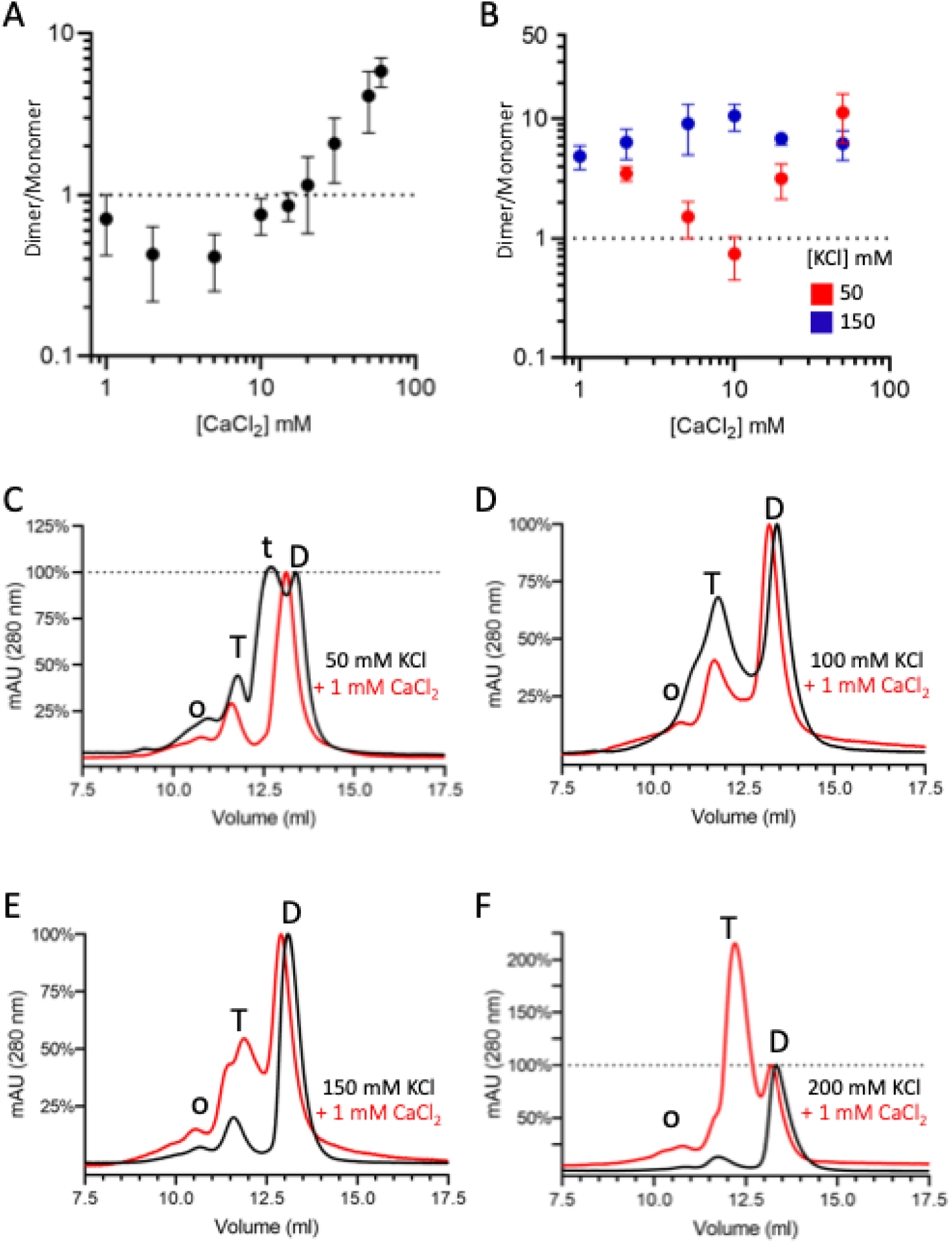
Electrostatic forms of CASQ2 assemblies compete with Ca^2+^-dependent dimerization and tetramerization. **A)** Dimer-to-monomer ratio of 10 nM CASQ2 as a function of increasing Ca²⁺ concentrations, as measured by mass photometry (MP). A slight decrease in dimer abundance is observed in the 1–5 mM Ca²⁺ range, followed by a marked increase at concentrations above 10 mM, indicating a threshold for Ca²⁺-dependent dimer stabilization. Data are presented as mean ± SD from at least two independent measurements performed with the same sample preparation, diluted in the same buffer. **B)** Dimer-to-monomer ratio of 25 nM CASQ2 as a function of increasing Ca²⁺ concentrations, in 50 mM and 200 mM KCl, as measured by mass photometry (MP). A decrease in dimer abundance is observed in the 10 mM Ca²⁺ range, followed by a marked increase at higher concentrations of Ca²⁺. Data are presented as mean ± SD from at least two independent sample preparations. **C-F)** SEC chromatograms of CASQ2 injected at 2 mg/ml in buffer containing either 50 mM, 100 mM, 150 mM, or 200 mM KCl (black chromatograms). After 1 h incubation in each ionic condition with 1 mM CaCl_2_, the sample was injected at 2 mg/ml into the SEC running in the respective buffer (containing either 50 mM, 100 mM, 150 mM, or 200 mM KCl and 1 mM CaCl_2_, red chromatograms). The curves were normalized to the height of the peak of the dimer to facilitate comparison across species: dimer (D), trimer (t), tetramer (T), and oligomer (o).

### A complex interplay between electrostatic and Ca^2+^-specific effects shapes CASQ2 polymerization

To dissect the role of electrostatics on the kinetics of Ca^2+^-dependent CASQ2 polymerization, we measured turbidity end-point values after 40 minutes since CASQ2 exposure to the physiological 1 mM free Ca^2+^ concentration^32^ across a range of ionic strengths (Figure 5A). The first prominent effect of K^+^ is a biphasic contribution to the maximal turbidity induced by Ca^2+^, in line with the previously observed consequences of increasing CASQ2 surface charge neutralization: progressive neutralization of intra-monomer and inter-monomer charge repulsion (promoting tertiary folding and oligomerization), up to a point where excessive charge screening and thickening of the solvation shell impede inter-particle interactions. The cooperative effect of K⁺ on CASQ2’s Ca^2+^ polymerization was additionally tested by comparison of the CASQ2 polymerization sensitivity to varying [Ca^2+^] in 10 and 50 mM KCl (Figure 5B). In 50 mM KCl, lower Ca²⁺ concentrations (0.1 mM Ca^2+^) are sufficient to induce a detectable turbidity increase than in 10 mM KCl (0.4 mM Ca^2+^). The 0.3 mM difference matches the Ca^2+^ levels needed for CASQ2 folding under low ionic strength conditions^10,16,27–31^, supporting that 50 mM K^+^ facilitates Ca^2+^- polymerization, at least in part, by promoting conformational compaction (Figure S2). On the other hand, the turbidimetric response to [Ca^2+^] >5 mM, a regime that also exhibits a pronounced Ca^2+^-dependent decline in turbidity (Figure 5A), is reduced at 50 mM KCl Vs. 10 mM KCl. This convergence also on the negative effect on Ca²⁺-dependent polymerization suggests that both K⁺ and Ca²⁺ contribute to excessive electrostatic shielding of CASQ2.

**Figure 5.**
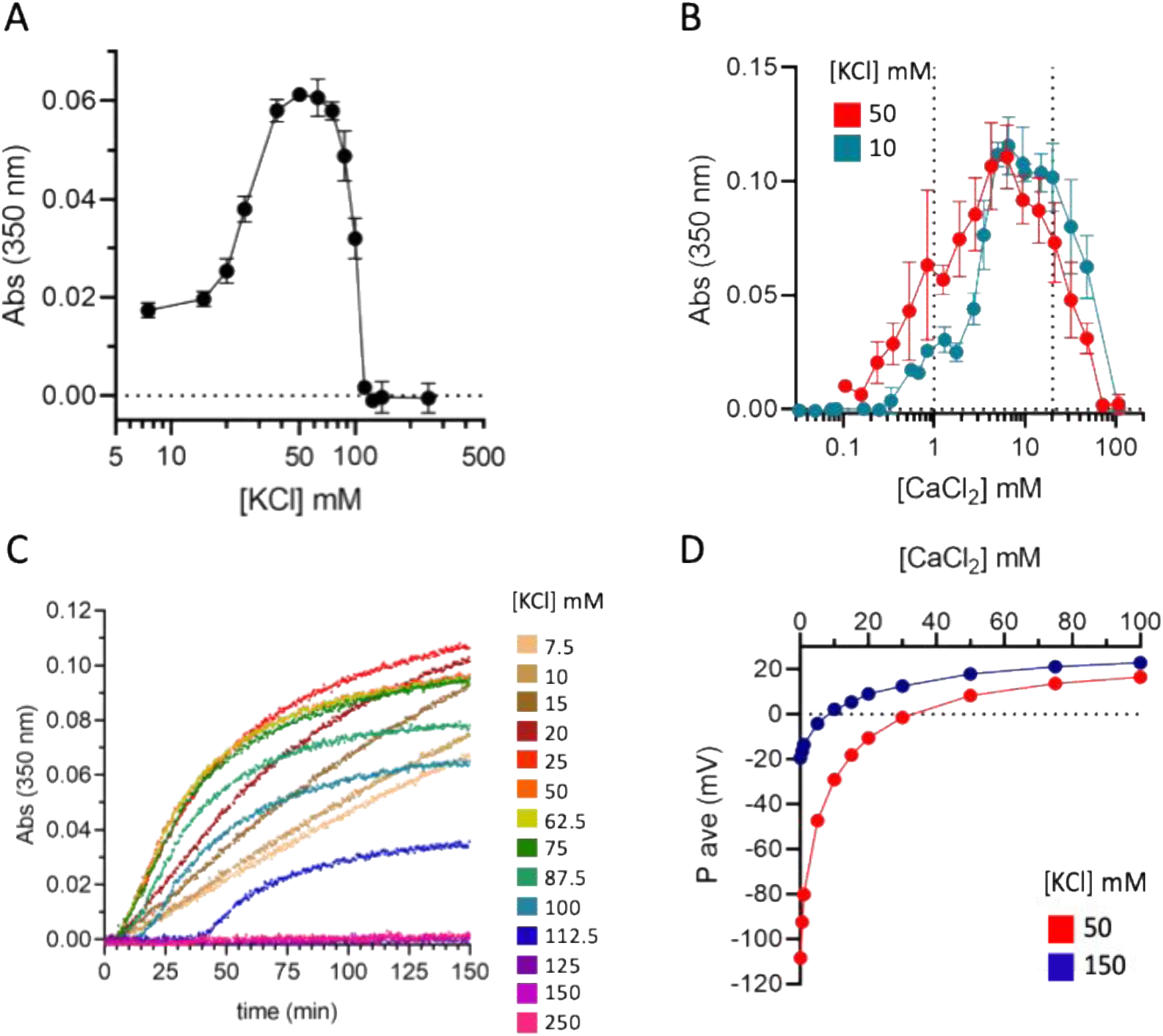
The Ca²⁺-specific polymerization of CASQ2 is a non-linear, electrostatically modulated process. **A)** End-point turbidity of CASQ2 after 40 minutes since exposure to 1 mM CaCl_2_ in varying ionic conditions (from 7.5 to 250 mM KCl). Data are mean ± SD from at least three independent experiments. **B)** Turbidimetric end-point measurements of CASQ2 samples after 40 minutes since the exposure to varying concentrations of Ca²⁺ (0.01–100 mM) in buffers containing 10 mM (pink) and 50 mM (purple) KCl. Data are presented as mean ± SD from three independent experiments. The estimates for the physiological concentration of free (1 mM) and total (20 mM) Ca^2+^ ions in the jSR^32^ are indicated by dotted lines. **C)** Turbidity assays showing sigmoidal growth kinetics of CASQ2 in response to increasing Ca²⁺ concentrations under different KCl conditions. Data are mean from three independent experiments. Data are shown as single point measurements from three independent experiments. **D)** ζ-potential (expressed in mV) of CASQ2 as a function of CaCl_2_ concentration. Of note, Ca²⁺ is markedly more efficient than K⁺ at neutralizing CASQ2 charges: To reach a 0 mV ζ-potential, 30 mM Ca²⁺ is needed in 50 mM KCl, versus only 3 mM in 150 mM KCl. Moreover, 10 mM Ca²⁺ is sufficient to compensate for the ∼75 mV difference brought by the 100 mM K^+^ difference between the two salt conditions.

To dissect more in depth the positive/ negative electrostatic contributions to CASQ2 Ca^2+^-dependent polymerization, we analyzed the kinetics of polymer formation (Figure 5C). A sigmoidal growth model fits all measurable turbidity curves ([KCl] <112.5 mM, with an average R^2^ = 0.986 ± 0.013, Figure 5A), from which it emerges that not only the amount of the soluble high-order polymers (deduced from the observed plateau, Figure S9A), but also the kinetics of polymerization is modulated by electrostatics. The nucleation phase is largely extended when [K^+^]>75 mM (Figure S9B), indicating that polymer nucleation does not rely simply on CASQ2 conformational compaction and/or destabilization of the competitor electrostatic oligomers, which instead benefit from K⁺ concentrations increasing up to 200 mM (Figures 2 and 4). Strikingly, the ionic condition beyond which nucleation is prolonged (∼80 mM KCl, 1 mM CaCl_2_; Figure S9B) corresponds to the switch point beyond which also the plateau phase is reduced (Figure S9A). Not only, the ζ-potential of this critical condition, estimated to fall between −45 and -50 mV, closely matches also that of the inflection point in maximal end-point turbidity measured for varying CaCl_2_ in 50 mM KCl (−45.7 ± 4.5 mV at 5 mM CaCl₂; Figure 5D). The convergence to a unique ζ-potential value, suggests that long-range electrostatics dominates the negative effect on both nucleation and the stability of CASQ2 Ca^2+^-dependent high-order polymers observable by turbidimetry. In turn, this raises the hypothesis that nucleation implies specific Ca²⁺-sensitive quaternary states, which formation is not linearly dependent on dismantling of non-specific electrostatic assemblies (as they would otherwise be supported by 75-200 mM KCl conditions, Figure 4), but which are sensitive *per se* to ionic conditions.

Adding a further layer of complexity, the same critical electrostatic condition defining the boundary between a positive and a negative contribution of K^+^ on the nucleation efficiency and solution stability of Ca^2+^-dependent CASQ2 polymers, also falls at the transition point from a poorly cooperative regime of polymer growth (Hill slope between 1 and 2, Figure S9C) to a highly cooperative one (Hill slope >2). The increase in the productivity of CASQ2 interaction beyond 75 mM KCl suggests the positive contribution of the protein’s conformational compaction and the parallel disassembly of otherwise competitive electrostatic interactions (Figures 2 and 4).

Despite their complexity, these observations are fully consistent with published end-point turbidimetric data (Figure S10) ^7,11,12,14,17,26,27,35^. Across studies, reducing monovalent cations enhances both the sensitivity and the amplitude of CASQ2 response to Ca²⁺. The fact that no positive effect of monovalent ions has been identified is due to the fact that previous studies did not explore the critical ζ-potential window (below –70 mV, corresponding to 20 mM NaCl or 50 mM KCl), where K⁺ ions exert a positive effect on Ca²⁺-dependent CASQ2 polymerization. Overall, our data support a model in which Ca²⁺-dependent CASQ2 polymerization is governed by a balance between positive and negative electrostatic effects, where Ca^2+^ and K^+^ can either cooperate positively or negatively.

Notably, while establishing electrostatic tuning as a central principle governing the Ca²⁺-dependent assembly of CASQ2, our data do not exclude additional ion-specific conformational effects on CASQ2 self-assembly properties. As a possible hint to this, what is intriguing is the small hinge in the CASQ2 polymerization end-points measured between 0.8 and 2 mM CaCl₂ (Figure 5B), consistently reproduced in each single experimental replicate in 10 mM KCl (Figure S11). A similar bi-phasic behavior, at the same Ca^2+^ concentrations, was independently reported as a variation in the Ca^2+^-binding affinities of recombinant CASQ1 and CASQ2 proteins^6^. Hence, mirroring the existence of distinct Ca^2+^-binding modes^6^, the biphasic Ca^2+^-dependent turbidity increase here observed may indicate a transition between distinct polymerization modes. This phenomenon does not relate to long-range electrostatic effects, as it alters the slope of polymerization (Figure S11), and hence the cooperativity of CASQ2 self-assembly.

### CASQ2 polymerization is an ion-sensitive, Ca^2+^-triggered switch

We directly inspected the dimensions of the soluble intermediate CASQ2 assemblies during Ca^2+^-dependent polymerization, via a particle-size distribution (PSD) granulometric analysis of CASQ2 in 50 and 150 mM KCl, exposed to gradually increasing free Ca^2+^ concentrations. To this aim, we employed a particular Dynamic Light Scattering (DLS) method which allows measurement of turbid samples. Unexpectedly, intensity-weighted PSD measurements revealed that CASQ2 assembly does not proceed through a gradual, stepwise increase in particle size. Instead, once a critical Ca²⁺ concentration is reached, CASQ2 undergoes a sharp, highly cooperative transition between two distinct quaternary states (Figure 6A): i) the Ca^2+^-independent oligomeric state with particles of 6-12 nm in diameter, in line with previous SEC and SAXS experiments, and consistent with dimers, tetramers, and possibly octamers; and ii) the Ca^2+^-induced high-order polymeric states, hundreds of nanometers large. Notably, even under 150 mM KCl conditions, which do not produce any measurable increase in turbidity (Figure 5C), large CASQ2 assemblies are formed. The lack of a corresponding turbidimetric signal likely reflects the formation of assemblies that are reduced in either number, size, and/or refractive index (presumably more hydrated). Overall, these data reveal for the first time that CASQ2 operates as a Ca^2+^-sensitive switch, oscillating with high cooperativity between an oligomeric phase and a high-order polymeric phase.

**Figure 6.**
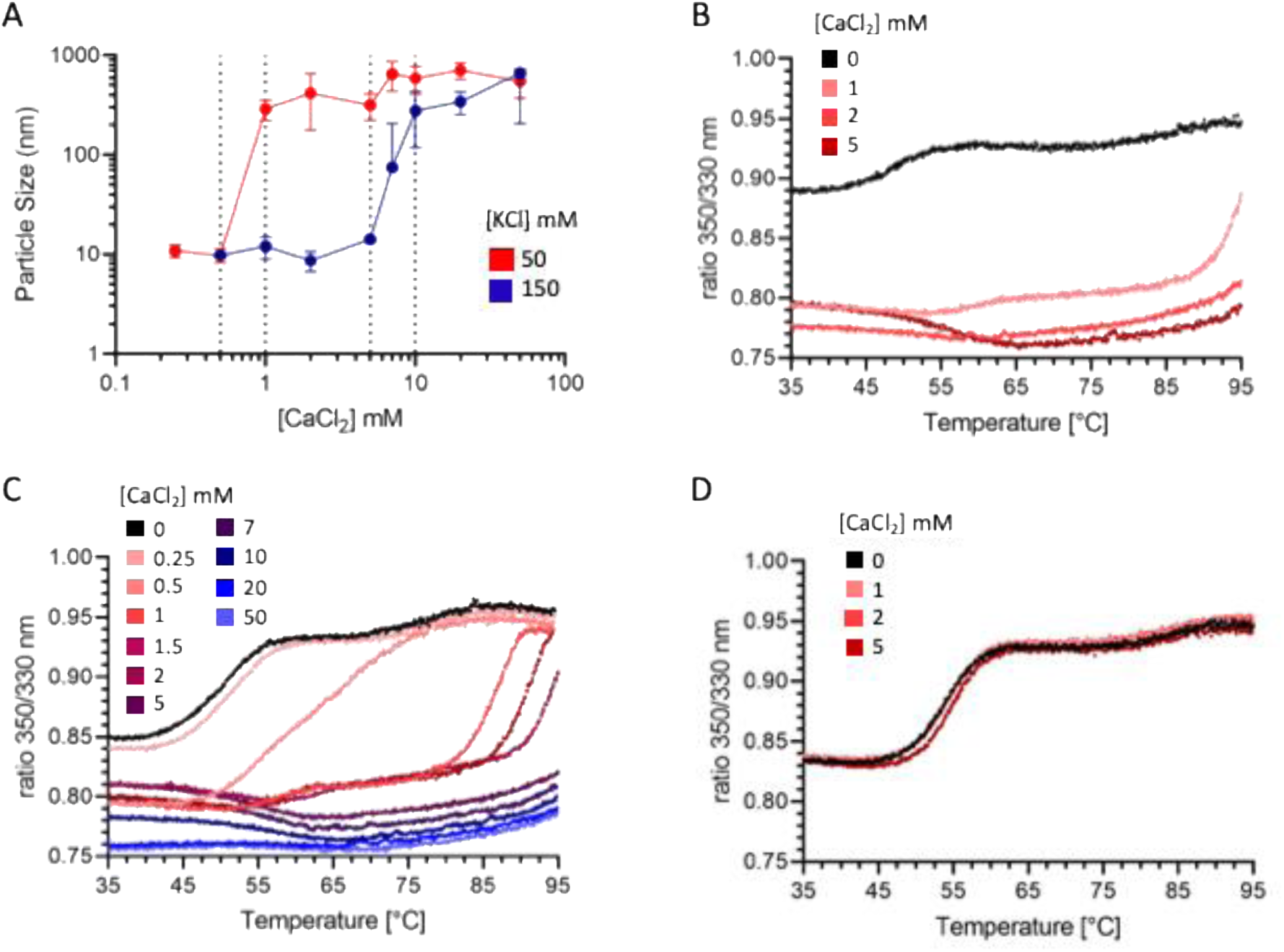
CASQ2 is a Ca^2+^-sensor oscillating between an oligomeric and a high-order polymeric state. **A)** Average size of CASQ2 particles in solutions of varying Ca²⁺ and K^+^ concentrations, as determined by Dynamic Light Scattering (DLS). In 50 mM KCl, CASQ2 oligomeric particles grow in size in response to 0.5 mM and 1 mM Ca²⁺, indicating polymerization. In the presence of 150 mM KCl, higher Ca²⁺ concentrations (5 to 10 mM) are required to promote particle size increase. Data are presented as mean ± SD from at least three independent measurements. **B-D)** Thermal denaturation curves, monitored by intrinsic tryptophan fluorescence (350/320 nm ratio), for CASQ2 samples in varying CaCl_2_ concentrations, and either 50 mM KCl (B), 150 mM KCl (C), or 300 mM KCl (D). Data are presented as mean from three independent measurements.

This switch-like behavior was confirmed by spectrophotometric and fluorimetric analyses in either bulk solution (cuvette) and glass capillaries. In cuvette, exposure of 1 mL solution of 2.5 μM CASQ2 to 50 mM CaCl_2_ triggers the formation, within seconds, of a separated protein-rich phase (Figure S12). At the same time, the thermal denaturation of CASQ2 in 50 mM KCl is enormously inhibited by addition of 1 mM CaCl_2_, up to 90°C (Figure 6B). As in PSD granulometry, this transition occurs over a narrow Ca²⁺ concentration range in both 50 mM and 150 mM KCl (Figures 6B, 6C). Increasing [KCl] suppresses this effect: the T_m_ of CASQ2 in 1 mM CaCl₂ is 8°C lower in 150 mM KCl than in 50 mM KCl (Figures 6C, S13), whereas any thermal stability shift is completely abolished at 300-500 mM KCl (Figure 6D). Overall, the sharp changes in the biophysical properties of CASQ2 in response to a narrow range of Ca^2+^ concentrations likely reflect the involvement of multiple packing interfaces and strongly cooperative formation of a highly cross-linked polymer.

### Limitations

In cardiac myocytes CASQ2 is strongly glycosylated during transport to the jSR^7^ and carries 0 to 2 phosphates per polypeptide^3,35^, whereas the recombinant CASQ2 protein used here does not carry post-translational changes. These may alter the balance of the specific effects of K^+^ and Ca^2+^ relative to those described herein, and/or the specific shape of the Ca^2+^-dependent polymers. In addition, while the estimated physiological concentration of CASQ2 falls within the 1-4 mM range^36–38^, the biochemical and biophysical properties of CASQ2 could be analyzed at much lower concentrations (with the exception of the MST data shown in Figure S7). Given the relevance of CASQ2 concentrations for its Ca^2+^-free oligomerization properties, this parameter should be taken into consideration when extending our findings to the physiological setting. One final limitation is that all experiments herein were performed in the absence of Mg^2+^. The effects of Mg^2+^ on CASQ2 Ca^2+^-dependent polymerization, due to their complexity, are beyond the current study and will be the subject of future research.

## Discussion

Our integrated experimental investigation redefines the molecular mechanism of cardiac CASQ2 polymerization by establishing three foundational advances: i) CASQ2 is physiologically a dimer; ii) Ca^2+^-dependent polymerization features a strong electrostatic component, and iii) CASQ2 polymerization is characterized by a switch-like transition.

Our results show for the first time that, under physiological ionic conditions, CASQ2 is intrinsically dimeric even at nanomolar concentrations and in the absence of its canonical ligand Ca^2+^. The relative ionic stability of this dimer with respect to other electrostatic assemblies hints at a structure match with the crystallographic dimer, where a network of hydrogen-bonds stabilizes the N-terminal tail swapping between monomers (Asp32 with Lys68 and Lys64; Lys68 with Glu74 and Glu78, Figure 1B)^8^; Additionally, this dimer appears to be functionally involved in sustaining the cooperativity of Ca^2+^-dependent polymerization (Figure 4), ensuring rapid responsiveness to physiological Ca^2+^ fluctuations during excitation-contraction coupling.

From SEC, MP, and turbidimetric kinetics (Figures 3 and 4), it appears that physiological amounts of K^+^ ions modulate the equilibrium between functional and non-functional CASQ2 assemblies: at concentrations below 50 mM, K^+^ enhances CASQ2 sensitivity to Ca^2+^ and the efficacy of Ca^2+^-dependent polymerization, likely by supporting protein folding and reducing inter-particle repulsion (Figures 5A, 5B); As the ionic strength increases and ζ-potential values approach 0 mV, unspecific inter-particle sticking (Figures 3A, 4A, 4B) is destabilized and the equilibrium shifts toward a non-electrostatically dependent, Ca^2+^-free CASQ2 dimer, competent for Ca^2+^-dependent quaternary assembly (Figure 4F). This form of dimer is expectedly a protagonist of the high cooperativity of Ca^2+^-polymerization response (Figures 5C, S9B), yet it is not the only player in the process of polymer nucleation (Figures 5B-C, S9A, S9C, 6A) and stabilization of the high-order Ca^2+^-dependent polymers (Figures 5A, 6D), which are ion-sensitive processes. Overall, the rapidity and magnitude of CASQ2 response to Ca^2+^ is optimal within a narrow electrostatic window between −60 and −40 mV, suggesting long-range interactions between CASQ2 particles, presumably involving a conformational switch of CASQ2 surface properties, tunes CASQ2 responsiveness during cyclic Ca²⁺ release in the jSR (Figure 6A). These experimental conclusions are independently corroborated by a theoretical model (Supplementary Text 1), which treats CASQ2 as a charge-regulated ion-binding electrolyte. This zero-dimensional electrostatic model simulates with impressive accuracy the interplay between electrostatic forces and the specific CASQ2 responsiveness to Ca^2+^, confirming that the interaction between Ca^2+^ ions and CASQ2 is characterized by a strong electrostatic component.

Finally, our data also reveal that CASQ2 does not polymerize through a linear assembly, exclusively Ca^2+^-driven, of monomers into dimers, tetramers and beyond. Instead, polymerization is the result of a Ca^2+^-triggered switch, oscillating from a dimeric/oligomeric pool to a structurally distinct polymeric state. This transition is highly cooperative, as shown by granulometry, thermal denaturation, and turbidity, and is sensitive to both ionic and protein concentrations. These experimental data and theoretical model provide the long-awaited experimental evidence required to build a model for CASQ2 polymerization, and for the interpretation of the many missense mutations spread all over the surface of cardiac CASQ2 in CPVT2. Specifically, the existence of an equilibrium between non-functional and functional dimers, and the ability of CASQ2 to dimerize at low nanomolar concentrations and in physiological ionic conditions, carry important physio-pathological implications: CASQ2 shall dimerize readily after translation and before its transport to the jSR, even in the absence of Ca^2+^. Building on the fact that CASQ2 trafficking is regulated by its quaternary assembly properties ^5,35,39^, and on published structural analysis of the missense mutations of CASQ1/2 ^3^, we propose that the dimerization competence of a mutant defines two distinct trafficking routes for either the dominant or recessive pathological mutants. While recessive mutations, affecting the stability of either the monomer or the dimeric state of CASQ2, are more easily recognized and not trafficked to the jSR or rapidly recycled from it, dominant negative mutations instead, by not altering the protein’s dimerization, are not recognized as non-wild-type and thus trafficked to, or maintained into, the jSR. As a consequence, the dominant negative CASQ2 mutants, residing in the same biological compartment with the wild-type counterpart, sequester it into defective polymeric structures, ultimately leading to Ca^2+^-leakage from the jSR and cardiac arrhythmias (Figure 7). This hypothesis fits with the fact that all published animal models of the recessive form of the CASQ2-related disease ^18,40–43^ feature a major loss of CASQ2 protein residing in the jSR, and a lower SR Ca^2+^ content, whereas the only dominant model for a CASQ2-related pathology ^44^ features a defect in the dynamics of SR Ca^2+^ exchange, but no evident alteration of either the content of CASQ2 or Ca^2+^ in the jSR.

**Figure 7.**
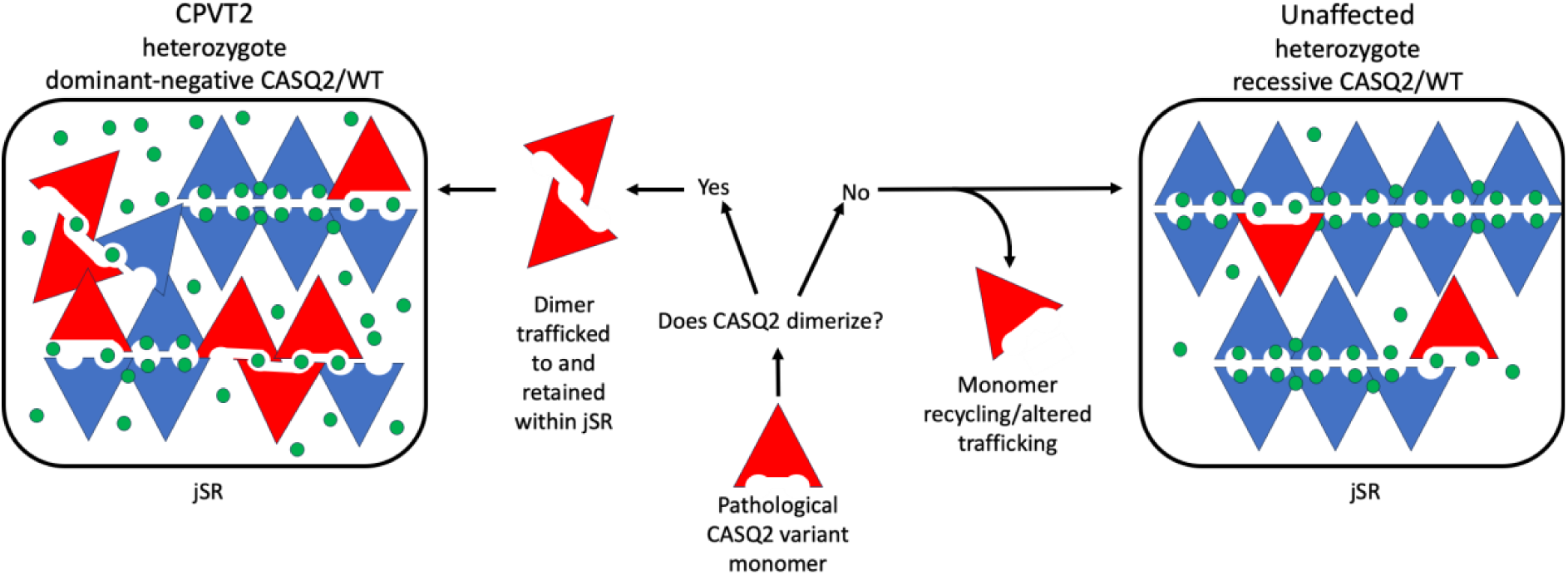
The proposed hypothesis on the distinct molecular pathways of recessive and dominant negative CASQ2 variants. Recessive CASQ2 mutations, affecting the stability of the protein monomer/dimer, and thus impeding necessary interactions for trafficking and retaining within the jSR, do not allow residence of the pathological protein variant within the jSR. For recessive CASQ2 mutations in heterozygosis, the wild-type counterpart can counteract the missing protein. Dominant negative CASQ2 variants, escaping the checkpoint for jSR delivery and/or residence, sequester the wild-type protein present within the jSR into a less-functional polymeric form. An altered hetero-polymeric mutant/WT-CASQ2-Ca^2+^ polymer may justify the slowly buffered, fast rise of free Ca^2+^-ions in the jSR detected for the dominant negative CASQ2 variant K180R ^44^ and in turn the inappropriate opening of the RyR2 channels and the predisposition to arrhythmias. CASQ2 pathological variant is in red. WT CASQ2 is in blue. Calcium ions are green.

## Materials and Methods

### Expression and Purification of cardiac Calsequestrin

The pET28a-based expression construct coding for N-terminally tagged His-CASQ2 [deleted of its first 19 residues] was transformed into the BL21 (DE3) E. coli strain. Overnight starter cultures were used to inoculate large-scale cultures (750 mL of broth per 2.8 L flask), which were grown at 37 °C to an optical density (OD) of 0.5 before induction with 1 mM IPTG. Following induction, cultures were incubated for an additional 3 hours at 30 °C. All cultures were grown in standard LB medium supplemented with 25 µg/mL kanamycin and 50 µg/mL chloramphenicol. Cells were harvested by centrifugation at 4,000 g for 20 minutes. The resulting cell pellets were collected and stored at -80 °C. Frozen cell pellets were resuspended in resuspension buffer (20 mM Potassium Phosphate, pH 7.4, 500 mM NaCl, 1 mM EDTA, 5 mM imidazole, 1 mM β-mercaptoethanol) and lysed mechanically using a Minilys® bead-beater (Bertin Technologies) at maximum speed (5000 rpm) for four cycles of 30 seconds each. The lysate was clarified by centrifugation at 8,900 × g for 1 hour at 4 °C. The supernatant was then filtered through a 0.45 μm membrane, and calsequestrin-containing fractions were isolated via immobilized metal affinity chromatography (IMAC) using a 5 mL HisTrap FF column on an ÄKTA go FPLC system. The IMAC buffers were as follows: Buffer A (20 mM KPi, pH 7.4, 300 mM NaCl, 20 mM imidazole) and Buffer B (20 mM KPi, pH 7.4, 300 mM NaCl, 300 mM imidazole). After a wash in 20% Buffer B, the protein was eluted in 100% Buffer B and subsequently dialyzed overnight at 4 °C in 2 L of Dialysis Buffer (20 mM HEPES, pH 7.3, 100 mM NaCl, 5 mM EDTA, 1 mM β-mercaptoethanol). Anion exchange chromatography was performed using a 5 mL HiTrap Capto Q column with Buffer A (20 mM HEPES, pH 7.3, 100 mM NaCl) and Buffer B (20 mM HEPES, pH 7.3, 1 M NaCl). Protein was eluted using a continuous gradient up to 100% Buffer B, with calsequestrin consistently eluting at 40–50% Buffer B. The calsequestrin-rich fractions were pooled and concentrated using an Amicon® centrifugal filter unit with a 10 kDa molecular weight cut-off. Size-exclusion chromatography (SEC) was performed by loading the sample into a 10 mL loop using a 5 mL syringe and allowing it to flow through a HiLoad 26/600 Superdex 200 pg column equilibrated with gel filtration (GF) buffer (20 mM HEPES, pH 7.3, 50 mM KCl). Calsequestrin-rich fractions were pooled, concentrated, aliquoted, and stored at -80 °C.

### Turbidity assays

The response of CASQ2 to CaCl₂ (0.01 mM to 100 mM) or MgCl₂ (0.01 mM to 100 mM) in the presence of 10 or 50 mM KCl was assessed using a Tecan Infinite 200® Pro plate reader by measuring absorbance at 350 nm. The assays were conducted in a 96-well plate, with each well containing a final volume of 140 µL. CASQ2 was buffer-exchanged to 20 mM HEPES (pH 7.3) and either 10 or 50 mM KCl. Prior to CaCl₂ or MgCl₂ addition, the plate containing CASQ2 sample was left at room temperature for 20 minutes and baseline turbidity was measured for blank determination. CaCl₂ or MgCl₂ was then manually added using a multichannel micropipette, and the sample was rapidly mixed by pipetting five times (avoiding the formation of air bubbles). Absorbance was recorded immediately after CaCl₂ or MgCl₂ addition. All measurements were performed at room temperature in triplicate for each condition.

### Mass photometry measurements

Mass photometry measurements were performed using a Refeyn TwoMP mass photometer (Refeyn Ltd). To prepare samples, CASQ2 was first buffer-exchanged into a measurement buffer containing 20 mM HEPES (pH 7.3) and a KCl concentration ranging from 25 mM to 500 mM. Each condition was incubated on ice for 45 minutes before being diluted in the equilibration condition to 100 nM CASQ2 in the first dilution and 10 nM CASQ2 in the final assessed drop. Data acquisition was initiated immediately after sample loading and resuspension by pipetting and recorded for 60 seconds. Molecules landing on the glass–buffer interface were visualized as changes in the interferometric scattering signal, which were automatically detected and quantified using AcquireMP software. To determine molecular mass, recorded contrast values for individual landing events were converted using a calibration curve established with protein standards. Data analysis was conducted using DiscoverMP software (Refeyn Ltd.), generating mass distribution histograms and applying Gaussian fits to determine the average molecular mass and sample heterogeneity. All experiments were performed at room temperature.

### Size-exclusion chromatography

A Superdex 200 10/300 GL column (Cytiva), connected to an ÄKTA go FPLC system, was equilibrated with a gel filtration (GF) buffer containing 20 mM HEPES (pH 7.3) and different concentrations of KCl (ranging from 50 mM to 200 mM) depending on the experimental condition being tested. A 100 µL sample of CASQ2 at 1 mg/mL, preincubated with the respective KCl concentrations, was injected into a 500 µL loop immediately after the addition of 1 mM CaCl₂. To evaluate the effect of CASQ2 concentration, the same column was used with a consistent GF buffer composition (20 mM HEPES, pH 7.3, 50 mM KCl) across all conditions. CASQ2 samples (100 µL) at varying concentrations (ranging from 5.6 µM to 225 µM) were injected, with 1 mM CaCl₂ added immediately prior to injection. All experiments were conducted at 4 °C, and elution profiles were monitored using the UV-Vis detection module integrated into the ÄKTA go system.

### Thermal stability assay

Protein samples were prepared in a buffer containing 20 mM HEPES (pH 7.3) with varying KCl concentrations (ranging from 25 mM to 500 mM) to achieve a final CASQ2 concentration of 2.5 µM. Approximately 10 µL of each sample was loaded into disposable Tycho capillaries (NanoTemper Technologies, cat# TY-C001). Thermal stability was assessed using the Tycho NT.6 instrument (NanoTemper Technologies). The system applied a temperature ramp from 35 °C to 95 °C at a controlled rate of 30 °C/min while recording the intrinsic tryptophan fluorescence at 330 nm and 350 nm. The fluorescence intensity ratio (350 nm/330 nm) was continuously monitored and used to generate an unfolding profile. Inflection temperatures (Ti), which indicate the temperature at which significant unfolding occurs, were determined automatically by the Tycho software. Each curve is the average of three independent measurements.

### Micro Scale Thermophoresis (MST)

MST experiments were carried out using a Monolith NT.115 instrument (NanoTemper Technologies) with premium Monolith capillaries. Protein labeling and binding assays were conducted using the Monolith His-Tag Labeling Kit RED-tris-NTA 2nd Generation (Cat. No. MO-L018) following the manufacturer’s instructions. The target protein was fluorescently labeled and diluted in PBS-T buffer. Prior to measurement, samples were incubated at room temperature for 30 minutes to allow binding equilibrium to be reached. The MST was performed at 25°C. Data analysis was conducted using MO.Affinity Analysis software.

### Dynamic light scattering and Z-potential measurements

Particle size analysis was performed using the NANOTRAC Flex (Microtrac) instrument, which employs a unique dynamic light scattering method with a flexible probe, in batch mode. CASQ2 samples were diluted in a buffer containing 20 mM HEPES (pH 7.3) and either 50 mM or 150 mM KCl to achieve a final protein concentration of 2.5 µM. Before each measurement, a blank reading was recorded using the buffer alone. Particle size was assessed using the same CASQ2 sample, with CaCl₂ added stepwise. The sample was continuously agitated by a piston before and during each measurement. At least six independent measurements were recorded for each CaCl₂ concentration. The Z-potential of CASQ2 at varying CaCl₂ and KCl concentrations was determined using a STABINO ZETA (Microtrac) ζ-potential analyzer. Similar to the particle size analysis, CASQ2 samples were diluted in a buffer containing 20 mM HEPES (pH 7.3) and varying KCl concentrations, ensuring a final protein concentration of 2.5 µM. For Z-potential measurements in the absence of CaCl₂, separate samples were used for each ionic strength condition. In contrast, to assess the effect of CaCl₂, measurements were performed stepwise on the same sample, recording changes in ζ-potential after each sequential addition of CaCl₂. A piston with a cutoff size of 200 nm was employed in this case.

## Funding information

This work was supported by Fondazione Telethon, Italy (grant number GMR24T1117).

## Notes

### Competing Interest Statement

The authors have declared no competing interest.

