## Supplementary Figures and Text for "Cardiac Calsequestrin is a Physiological Dimer that Polymerizes through a Ca²⁺-Triggered Cooperative Switch"

**SUPPLEMENTARY MATERIALS**

### Supplementary Figures

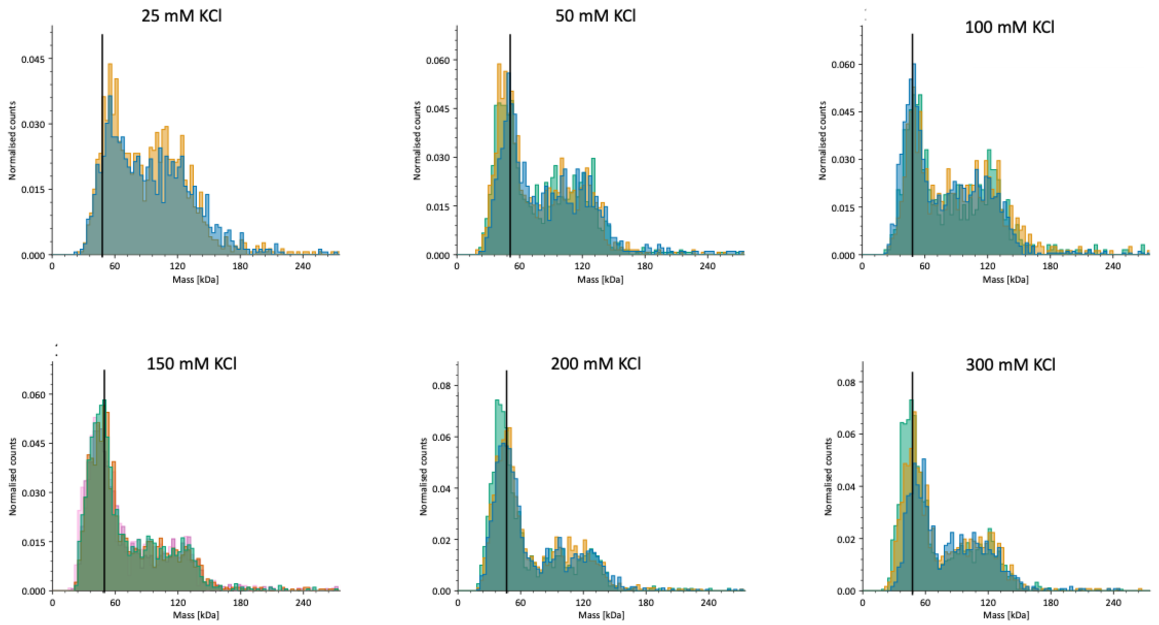


**Figure S1. Mass photometry (MP) of CASQ2 under increasing KCl concentrations.** MP measurements reveal the relative abundance and distribution of CASQ2 monomers, dimers, and higher-order oligomers in buffer conditions with increasing KCl concentrations (25–300 mM). Three independent measurements are shown for each condition. The 48 kDa apparent mass value is highlighted by a black vertical bar, to facilitate comparison of the apparent volume registered for the monomer species across ionic conditions. Of note, quaternary assemblies corresponding to dimers, trimers, and tetramers, form in absence of $Ca^{2+}$ and are progressively inhibited by increasing ionic conditions (in the 25 to 200 mM KCl range, see also Figure 3A). These results suggest a dynamic equilibrium between CASQ2 oligomeric states modulated by ionic strength, with $K^{+}$ ions influencing both the prevalence and stability of multimeric species.

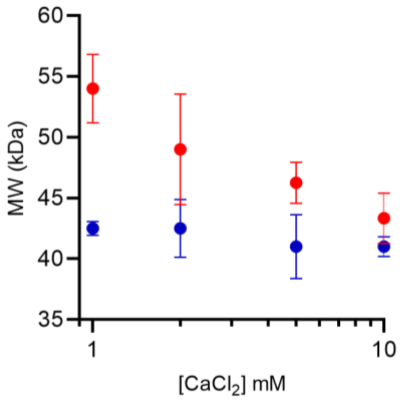


**Figure S2. Mass photometry (MP) analysis of the size of CASQ2 monomer under increasing $CaCl_2$ concentrations and in two distinct ionic strength conditions (50 mM and 200 mM KCl).**

The apparent volume occupied by CASQ2 monomers in 50 mM KCl solution (red) decreases with increasing $CaCl_2$. In 200 mM KCl (blue points), $CaCl_2$ does not compact any further the folding of CASQ2 polypeptide. At least three independent measurements, with CASQ2 25 nM, were considered for each condition.

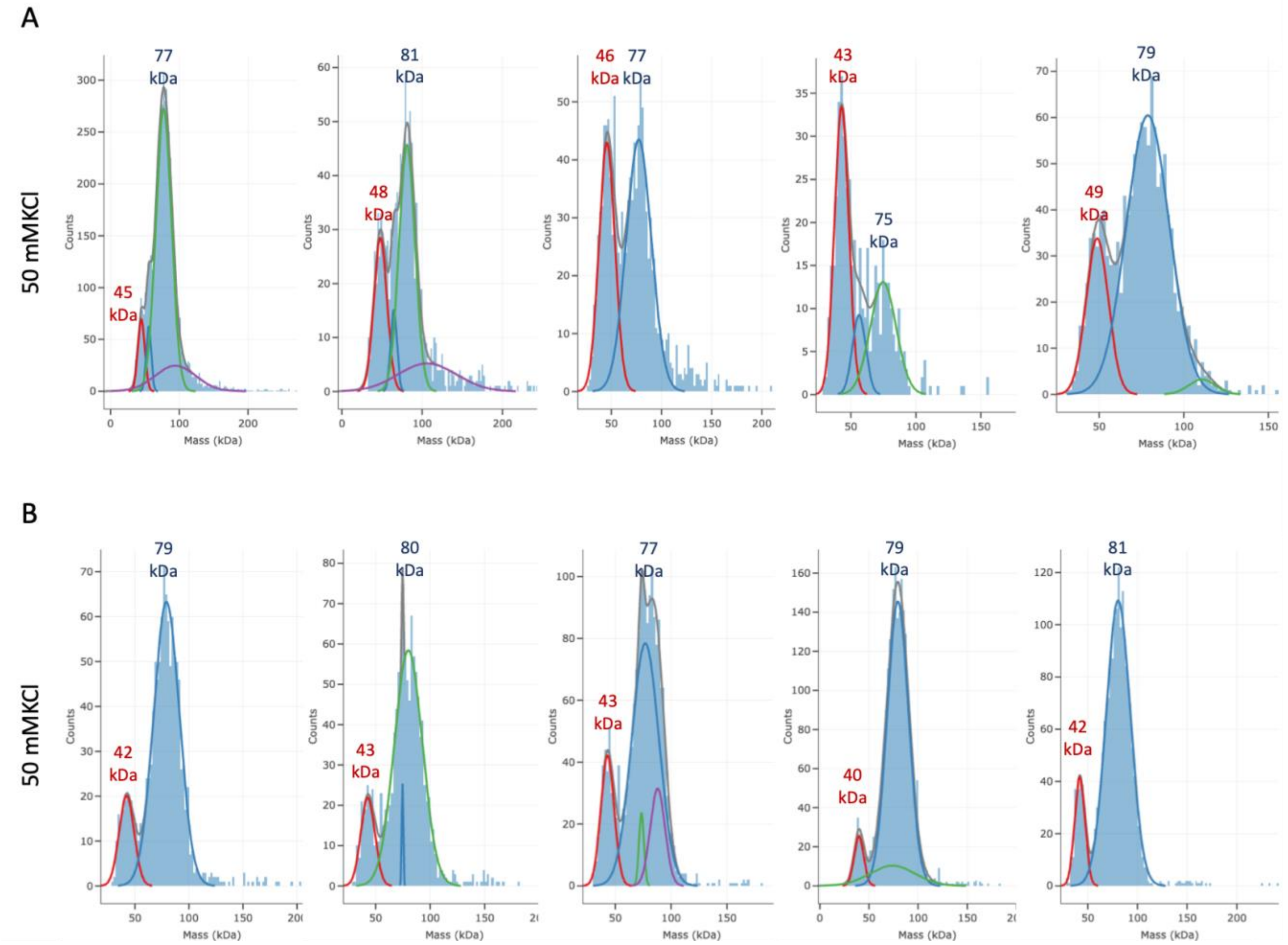


**Figure S3. Representative Mass photometry raw data of CASQ2 multimerization at increasing $Ca^{2+}$ concentrations.** Distributions of detected particle masses for 25 nM wild-type CASQ2 incubated with increasing concentrations of $CaCl_2$, reveals a shift from monomer-dimer peaks with increasing $Ca^{2+}$ levels. Data are fitted using automatic Gaussian distributions (PhotoMol [45]) to estimate peak centers and molecular mass assignments for the monomer (red) and dimer (blue) species.

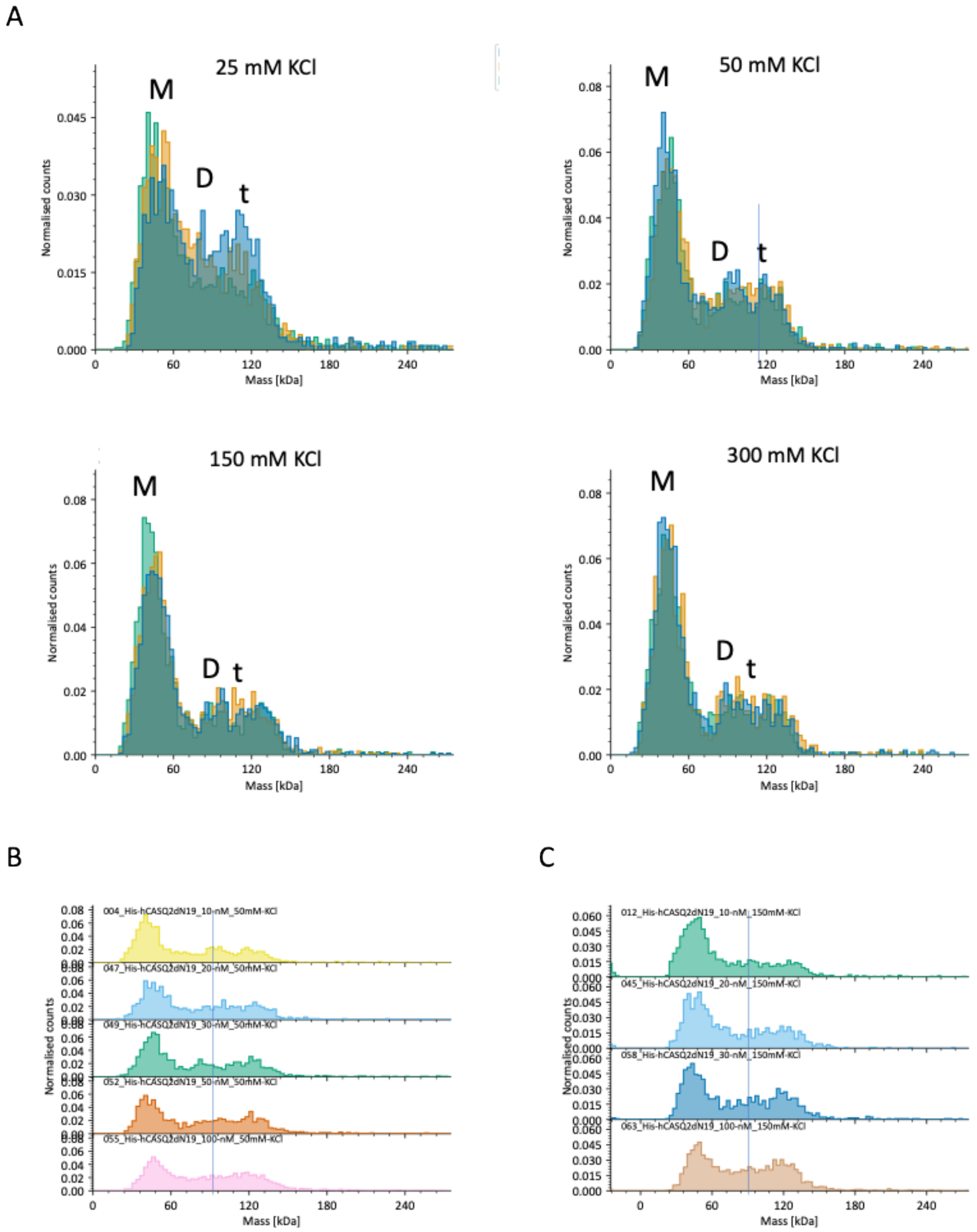


**Figure S4. MP analysis of CASQ2 oligomer distribution as a function of KCl concentration. A)** Normalized mass distributions for His-tagged CASQ2 at 10 mM protein concentration measured in 25, 50, 150, and 300 mM KCl. Mass distributions shift from broad, multimodal peaks at low salt toward narrower peaks centered around the size of dimeric (D) and trimeric (t) states, and a shoulder at the approximate mass of the tetramer (T), with increasing ionic strength. Moving from 25 to 50 and 150 mM KCl, a decrease in the proportion of the dimeric quaternary assemblies is evident. Above 150 mM KCl, no significant change in the proportion of the dimeric Vs. monomeric species is observed. Data show the overlay of the measurements of three independent samples. **B-C)** Replicate datasets for varying concentrations of CASQ2 (10, 20, 30, 50, and 100 nM) in 50 mM KCl buffer (B), and 200 mM KCl buffer (C). A vertical line positioned at the average mass of the dimeric species (80 kDa) is drawn as reference to facilitate visualization of the fact that supra-dimeric assemblies are sustained by increasing CASQ2 concentrations. These results support the interpretation that CASQ2 is able to form quaternary assemblies in absence of $Ca^{2+}$, also at low protein concentrations.

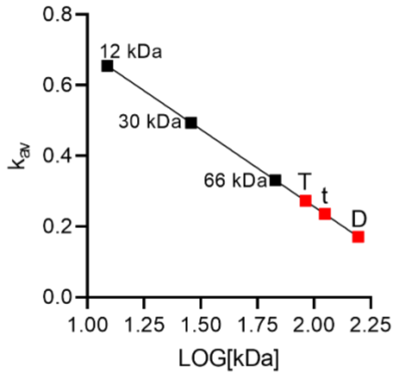


**Figure S5. Correlation of molecular weight and elution profile in SEC (Superdex 200 10/300) calibration curve.** Standard proteins of known molecular weight (black dots) were used to construct a calibration curve plotting the distribution coefficient (K_av) against the logarithm of their molecular weight ($\log_{10}$[kDa]). Red squares represent the estimated molecular weights of CASQ2 species based on their elution volumes: 90 kDa (dimer, D), 110 kDa (trimer, t), and 152 kDa (tetramer, T), interpolated on the standard curve. This calibration was used to assess the quaternary state of CASQ2 under different ionic conditions.

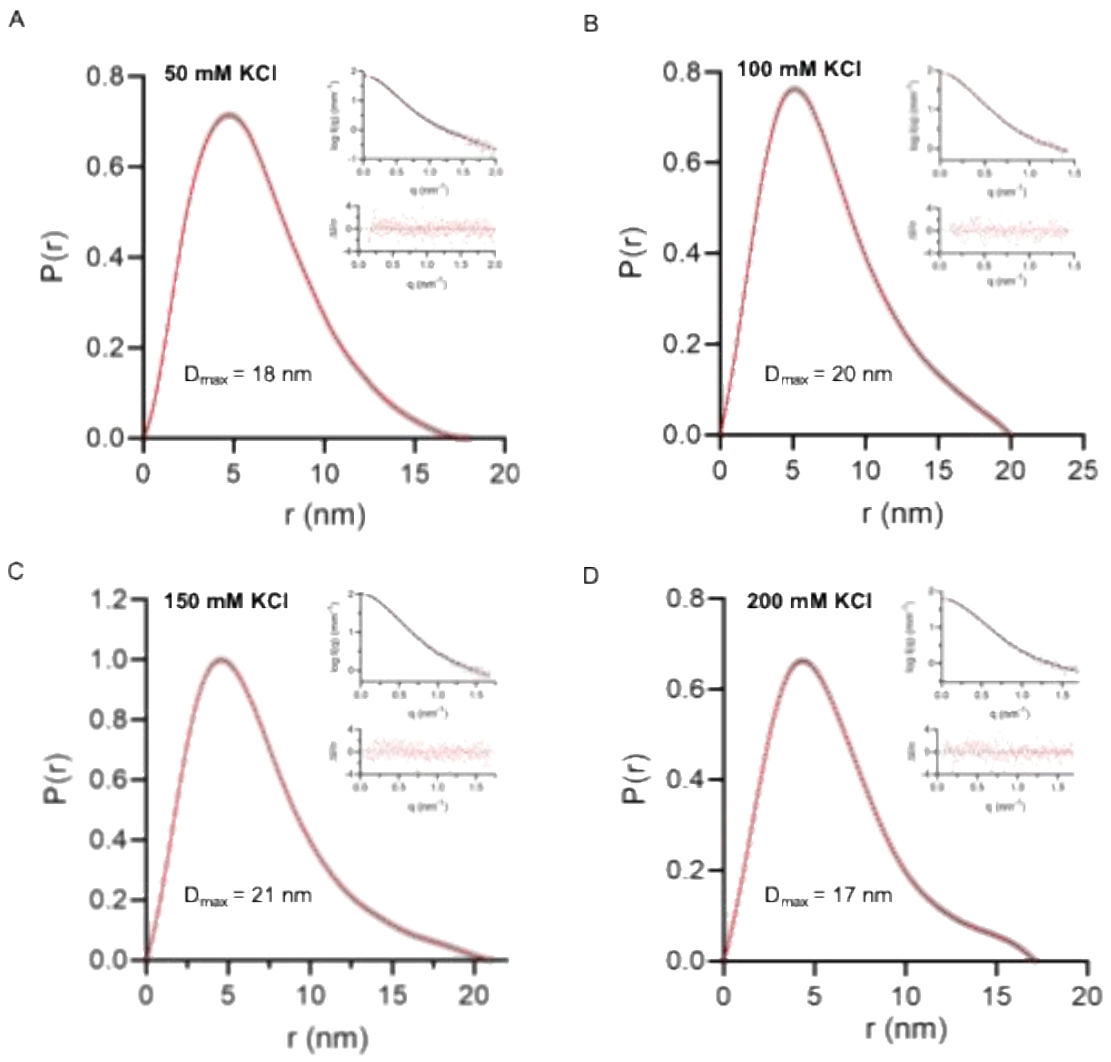


**Figure S6 – P(r) graphs for fixed 14 uM CASQ2**. Probability distribution (P(r)) functions of CASQ2 batch-SAXS experiments performed in increasing [KCl] (A-D). Insets show data used for the P(r) computation and fit (top) with the corresponding residuals (bottom).

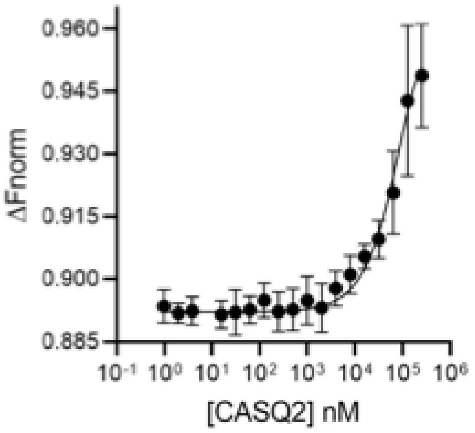


**Figure S7.** MST assay of CASQ2 self-association in PBS buffer, showing similar pronounced concentration-dependent CASQ2 self-association. Data are presented as mean ± SD from three independent measuremen

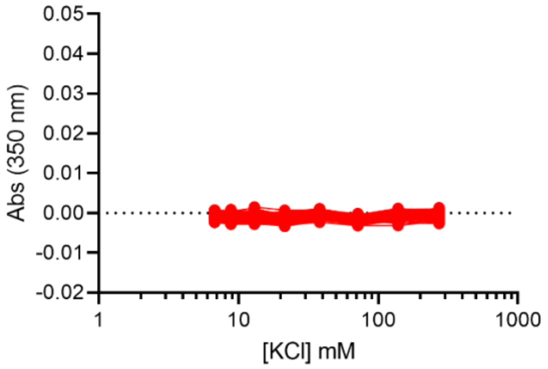


**Figure S8. Turbidimetric analysis of CASQ2 multimerization in the absence of divalent ions.** Absorbance at 350 nm was measured for 45 minutes since the incubation of CASQ2 with increasing concentrations of KCl, in the absence of $Ca^{2+}$ or $Mg^{2+}$. The lack of significant turbidity changes across the tested ionic strengths confirms that CASQ2 does not efficiently form large multimers without the presence of divalent cations. These results support the requirement of $Ca^{2+}$ or $Mg^{2+}$ to promote higher-order assembly of CASQ2.

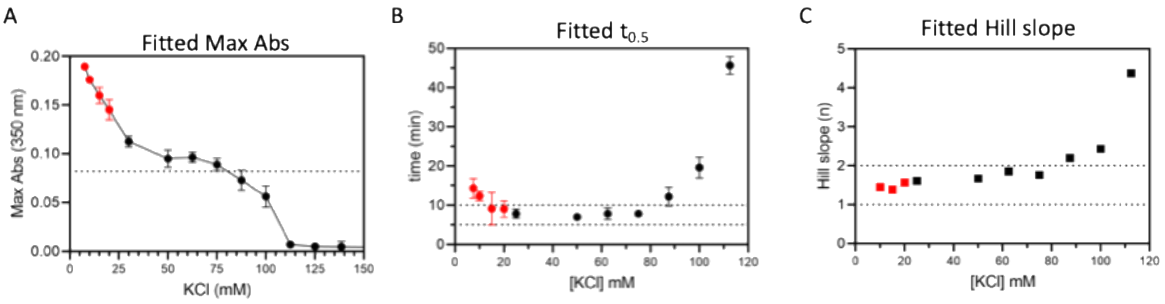


**Figure S9. Parameters calculated from the fitted sigmoidal growth curves for $Ca^{2+}$-dependent polymerization.** For [KCl] < 15 mM, the values are highlighted in red, as only the initial linear growth phases were captured, limiting the accuracy of the mathematical model for those conditions, especially for the maximal absorbance level reached at plateau. Of note, we experimentally observed an increased precipitation in time for those low-ionic conditions, which reflects the insoluble nature of the polymers formed. For each panel, values are reported as mean $\pm$ SD across triplicates. **A)** Maximal absorbance levels reached at plateau. The plateau phase, at which $Ca^{2+}$-CASQ2 polymers are at an equilibrium in solution, is similar across 50-75 mM [KCl]. For increasing KCl concentrations, the amount, and/or number, and/or extinction coefficient of the polymers at equilibrium is reduced. The dotted line defines the boundary between the supportive and the negative role of $K^+$ on the stabilization of high-order CASQ2/$Ca^{2+}$ soluble polymers. **B)** Hill slope values show positive cooperativity of polymerization at moderate ionic strengths (25–75 mM KCl), rapidly increasing above 100 mM KCl**.** The dotted lines define the boundaries of the poorly cooperative regime. **C)** The nucleation phase, defined as the time required to reach half the maximum turbidity, increases for [KCl] > 90 mM.

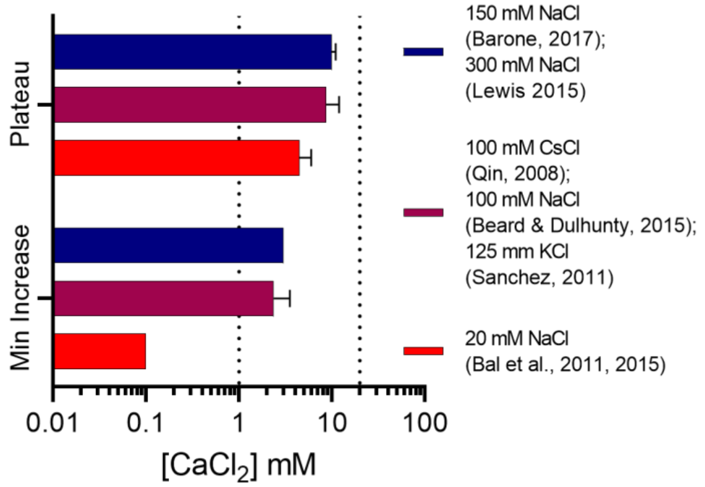


**Figure S10.** Overview of the $Ca^{2+}$-dependent CASQ2 polymerization main parameters extracted from published data: i) the minimal $Ca^{2+}$ concentrations required to elicit the onset of turbidimetric increase, and ii) the minimal $Ca^{2+}$ concentrations required to reach the the maximal responsiveness plateau phase. Data are divided according to the ionic strength of the published experiment, as reported in the legend. The estimates for the physiological concentration of 1 mM free and 20 mM total $Ca^{2+}$ ions in the junctional Sarcoplasmic reticulum (jSR) [32] are indicated by dotted lines. The ζ-potential of the lowest ionic strength condition reported in literature (20 mM NaCl, corresponding to -105.7 mV, Figure 2D) is matched by our 50 mM KCl condition (–108.4 mV, Figure 2D), which explains the fact that no positive effect is observed for $K^+$ on CASQ2 $Ca^{2+}$-dependent polymerization

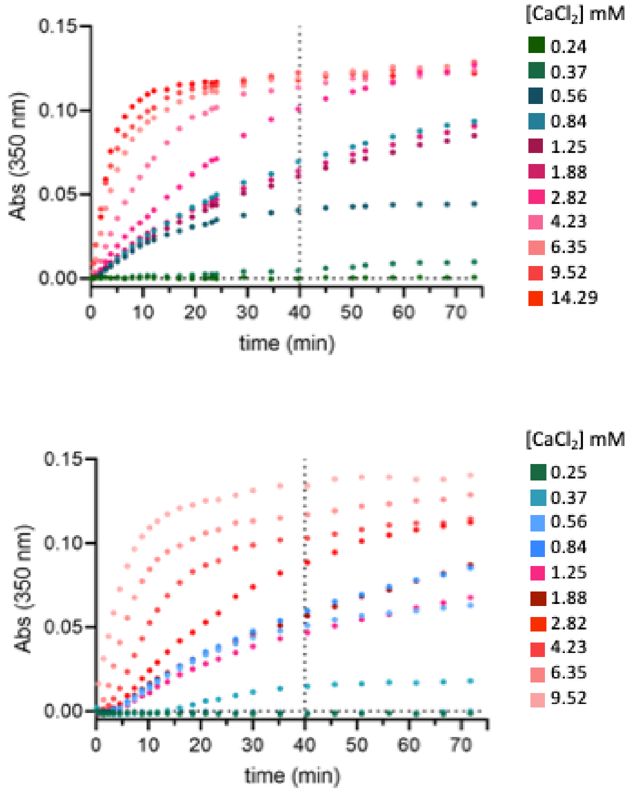


**Figure S11. Turbidity of CASQ2 protein in response to the addition of varying concentrations of $CaCl_2$**. For each experimental condition, the absorbance measurements are reported as the average of three measurements from three independent samples at defined timepoints. The legend on the right refers to the final concentration of $CaCl_2$ once added to the sample. In the range 0.5-1.25 mM $CaCl_2$, the initial turbidity slope is reproducibly higher than the response to 1.88 mM $CaCl_2$.

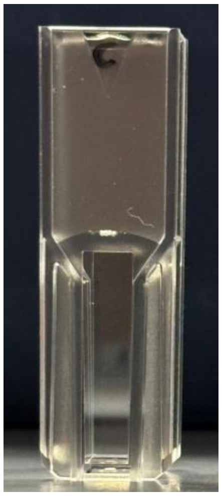

**Figure S12. CASQ2 polymerization at high $Ca^{2+}$ concentrations results in visible phase separation.** Representative image of a cuvette containing 2.5 µM CASQ2 in 50 mM KCl buffer exposed to 50 mM $CaCl_2$, showing a visible phase-separated system. The appearance of a dense, turbid layer indicates the formation of polymeric CASQ2 condensates, supporting the occurrence of a macroscopic phase transition upon calcium-induced polymerization. In addition, measurement of the absorbance at 280 nm of samples from all three liquid phases (above, within, and below the turbid phase) reveals that only in the turbid phase, an absorbance curve appropriate for a protein is measurable (corresponding to CASQ2 concentration of about 20 µM). This visual evidence complements turbidimetry, DLS, and thermal shift assays in demonstrating the sharp transition between oligomers and high-order CASQ2 polymers.

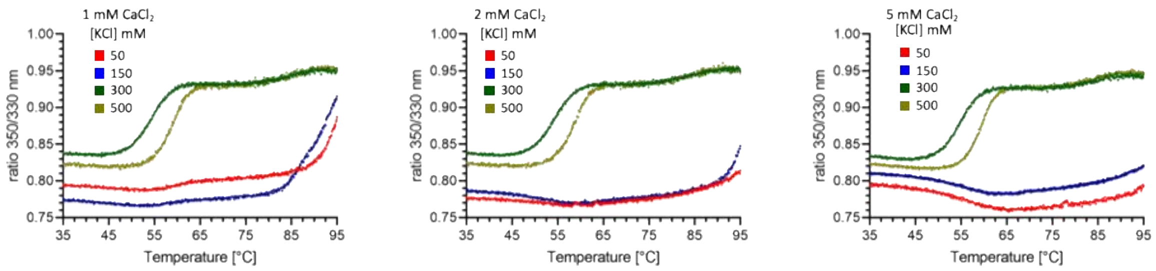


**Figure S13. Thermal stabilization of CASQ2 by $Ca^{2+}$ is progressively inhibited by increasing KCl concentrations.** Thermal denaturation profiles of CASQ2 (6 µM) after 1 h incubation with 1, 2, or 5 mM $CaCl_2$ in buffers containing 50, 150, 300, or 500 mM KCl. In 50 mM KCl, $Ca^{2+}$ induces a pronounced upward shift in melting temperature (Tm), consistent with formation of highly stabilized CASQ2 polymers. This $Ca^{2+}$-induced thermal stabilization is reduced in 150 mM KCl and abolished at 300–500 mM KCl, demonstrating that high concentrations of monovalent cations antagonize the formation of the $Ca^{2+}$-dependent polymeric state.

**Supplementary Text 1**

**A 0-D theoretical formulation describing the formation of dimeric and polymeric CASQ2 assemblies, based purely on electrostatics.**

**SCOPE:**

The following approach, while guided by physical intuition, is phenomenological. It is aimed at qualitatively accounting for the most important experimentally observed effects of different ionic environments (containing $K^+$, $Na^+$ and/or $Ca^{2+}$) on the **equilibrium self-association properties of wild-type human Calsequestrin 2 (CASQ2) without post-transcriptional modifications**.

No mathematical fitting of parameters has been attempted. All values used in this model have either been calculated or set by hand.

**PREMISES:**

The current theoretical scheme is built under the following premises:

(1) Depending on the ionic environment, CASQ2 molecules partition between 2 states: *optimally folded* molecules (capable of dimerization and $Ca^{2+}$-dependent polymerization, see next premise) and *non-functionally folded* ones (capable of forming electrostatic dimers, non-competent for $Ca^{2+}$-dependent polymerization). Electrostatically-driven oligomers, described in this manuscript, are neglected from this theoretical scheme for convenience.
(2) CASQ2 polymerization can only occur among optimally folded molecules
(3) CASQ2 dimerization can occur only between molecules in the same state, i.e. mixed dimers are not allowed.
(4) CASQ2 polymerization occurs by sequential addition of dimers.

**REACTION SCHEME:**

CASQ2 is modeled as following the set of reactions below. These reactions are treated separately for convenience, as they try to reflect different aspects of the CASQ2 behavior: dimerization vs. polymerization

$$D_{nf} \rightleftharpoons 2M_{nf} \rightleftharpoons 2M_o \rightleftharpoons D_o \qquad [1]$$

Where D refers to a dimer, M to a monomer, the subscript *nf* refers to non-functionally folded CASQ2 monomers/dimers and the subscript *o* refers to optimally folded monomers/dimers. Reaction [1] has 3 equilibrium constants, named *K1* (reaction $D_{nf} \rightleftharpoons 2M_{nf}$), *K2* ($2M_{nf} \rightleftharpoons 2M_o$) and *K3* ($2M_o \rightleftharpoons D_o$).

Sequential addition of $D_o$ dimers leads to oligomers and polymers. At this point, we impose a maximum number of $D_o$ molecules per polymer, *N*, due to the otherwise difficult tractability of the equations.

$$ND_o \rightleftharpoons (N\text{-}2)\, D_o + T_o \rightleftharpoons (N\text{-}3)\, D_o + H_o \ldots \rightleftharpoons P_o\,(N\text{-}1) + D_o \rightleftharpoons P_o\,(N) \qquad [2]$$

Note that in the current formulation we have not allowed for polymers to form by addition of oligomers (e.g. an octamer may be obtained by combination of a dimer and a hexamer, as modelled, or by combination of two tetramers). We did not allow this second form of combination, due to the complexity in tracking individual reactions when N becomes high, and the lack of certainty regarding the equilibrium constants.

The equilibrium constants for reaction [2] are assumed to take the form:

$$K_{n\ dimers} = K_{tetramerization} \cdot \alpha^{(n\text{-}2)} \qquad [3]$$

Where $n$ refers to the number of $D_o$ dimers involved in the reaction. For example, when $n = 2$ (i.e. tetramer formation), $K_{n=2}$ reduces to $K_{tetramerization}$. For the formation of a hexamer (n = 3), $K_{n=3} = K_{tetramerization} \cdot \alpha$, and so on until the maximum polymer forms, $K_{n=N} = K_{tetramerization} \cdot \alpha^{(N-2)}$. Note that α denotes the cooperativity of the process (i.e. positive when α > 1).

The total concentration of CASQ2 is given in terms of $[M_o]$, the equilibrium constants, α and N. Once $[M_o]$ is known, the fractions of all other species are calculated. We will return to the numeric value of the equilibrium constants $K1$, $K2$, $K3$, $K_{tetramerization}$ and α at a later stage. At this point in time, we first need to discuss what are the driving processes of CASQ2's optimal folding in the current theory.

**PROCESSES:**

***Process 1: The neutralization of CASQ2 net charge depends on ionic environment***

This process builds on the intuition that proteins cannot be optimally folded unless their charge is neutralized by counter-ions. We used the canonical human CASQ2 sequence (Uniprot ID O14958-1) to determine the CASQ2 net charge at pH 7.3 (the pH used in the experiments of the current manuscript). Charges for each individual amino-acid were calculated based on the pKa values of the amino acids [REFS 1-2], and are shown here in summarized form:

Amino-acid charges: {'A', 'R', 'N', 'D', 'C', 'E', 'Q', 'G', 'H', 'I', 'L', 'K', 'M', 'F', 'P', 'S', 'T', 'W', 'Y', 'V'}, [0, 1, 0, -1, 0, -1, 0, 0, 0.17, 0, 0, 1, 0, 0, 0, 0, 0, 0, 0, 0].

Note that histidine (H) was given a charge of 0.17 due to partial protonation at pH 7.3. The calculated net charge of CASQ2 was (-57.98).

Ions were considered to neutralize CASQ2 charges depending on their charge density relative to $K^+$. The following formula accounted for the amount of cationic charge available to neutralize CASQ2 in any given ionic environment containing $K^+$, $Na^+$ and/or $Ca^{2+}$:

$$\textit{cationic\_charge} = [K^+] \cdot (K\_r/K\_r) + [Na^+] \cdot (K\_r/Na\_r) + 2 \cdot [Ca^{2+}] \cdot (K\_r/Ca\_r) \quad [4]$$

where K_r is the ionic radius of $K^+$ (133 pm), Na_r is the ionic radius of $Na^+$ (95 pm) and Ca_r is the ionic radius of $Ca^{2+}$ (65 pm). CASQ2 neutralization was expressed as a percentual value of its net charge:

$$\text{Non-neutralized charges (\%)} = 100 \cdot (\text{CASQ2\_charge} + \textit{cationic\_charge}/3.35) / \text{abs(CASQ2\_charge)} \quad [5]$$

In other words, a non-neutralized CASQ2 (as if there were no ions around it) would render a (-100%) value in [6], whereas a fully neutralized one would render a 0% value. Note the denominator that divides *cationic_charge:* it was set at 3.35 to match the $[K^+]$ leading to zero ζ-potential at the experimental concentration of 194.2 mM (see Figure 2D within the manuscript and Footnote 1 within this appendix).

---

**Footnote 1**: Experiments indicate that, when CASQ2 dwells within an environment containing KCl alone, ζ-potential equals 0 when $[K^+] = 194.2$ mM. We can use equations [4] and [5] (setting non-neutralized charges = 0) to estimate the value of X, the denominator of *cationic_charge* in equation [5].

$$\textit{cationic\_charge} = [K^+] \cdot (K\_r/K\_r) = 194.2 \quad [4\text{-special}]$$

$$0 = 100 \cdot (\text{CASQ2\_charge} + \textit{cationic\_charge}/X) / \text{abs(CASQ2\_charge)} \quad [5\text{-special}]$$

Replacing equation [4-special] into equation [5-special] and rearranging, we get

$$X = \textit{cationic\_charge} / (-\text{CASQ2\_charge}) = 194.2 / (-(-57.98)) = 3.34943$$

Also note that, by applying an analogous reasoning, we can roughly estimate the maximum number of $Ca^{2+}$ moles that can be bound to a fully neutralized mole of CASQ2 molecules. We show the end-stage of the calculation

$$Ca^{2+}_{Max\text{-}bound} = (-\text{CASQ2\_charge}) \cdot (3.35/2) \cdot (Ca\_r / K\_r) = 57.98 \cdot 1.675 \cdot (65 / 133) = 47.45$$

The number agrees with the experimentally estimated maximum $Ca^{2+}$-binding capacity of CASQ2 of 40-50 ions per molecule [3].

**Process 2: the fraction of optimally folded CASQ2 is proportional to charge neutralization**

The process above, charge neutralization, was assumed to drive the optimal folding of CASQ2. To model this, we used the following ascending sigmoidal equation:

$$Promoted\_folding = ((1 + (EC50_{Folding} / \text{ionic\_strength})^{1.5}))^{(-1)} \quad [6]$$

Where $EC50_{Folding}$ was defined to be the amount of *cationic_charge* (formula [4]) needed for 50% of the CASQ2 charges to be neutralized. Applying formula [5] with Non-neutralized charges = (-50%), we obtain:

$$EC50_{Folding} = 3.35 \cdot ( (-0.5) \cdot \text{abs(CASQ2\_charge)} - \text{CASQ2\_charge} ) = 97.0991 \text{ mM}$$

On what concerns ionic strength, it was calculated as dependent on charge density (i.e. not all ions count the same for ionic strength purposes, even if they have the same charge). Once again, $K^+$ was set as reference

$$\text{ionic strength} = 0.5 \cdot \sum (K\_r / Ion\_r) \cdot [Ion] \cdot Ion\_\text{charge}^2 \quad [7]$$

**Process 3: Excessive ionic strength promotes non-functional foldings of CASQ2**

Beyond a certain amount (mM) of ionic strength, proteins leave their optimal functional configuration due to excessive charge-screening from hydration shells, hampering electrostatic (and $Ca^{2+}$-dependent) interactions. We modeled this process as a descending sigmoidal curve with an $EC50_{Loss}$ defined to be the amount of *cationic_charge* (formula [4]) needed to revert 50% of the CASQ2 charges. Applying formula [5] with Non-neutralized charges = (+50%), we obtain:

$$EC50_{Loss} = 3.35 \cdot ( 0.5 \cdot \text{abs(CASQ2\_charge)} - \text{CASQ2\_charge} ) = 443.3034 \text{ mM}$$

And the descending sigmoidal curve representing this process is

$$Loss = ((1 + (\text{ionic\_strength}/EC50_{Loss} )^{1.5}))^{(-1)} \quad [8]$$

**Process 4: $Ca^{2+}/K^+$ competition right-shifts $EC50_{Loss}$**

The current manuscript presents strong evidence of $K^+/Ca^{2+}$ discrimination in the polymerization reaction of CASQ2 (see reaction [2]), plus a competitive reaction between $K^+$ and $Ca^{2+}$ in favor of the latter which leads to the dissipation of non-functionally folded structures. In order to account for these effects, the model calculates the $Ca^{2+}$-dependent ionic strength relative to the total ionic strength provided by cations (each calculated by formula [7])

$$bivalent\_related\_folding = (\text{Ca\_is}) / (\text{ionic\_strength} - \text{Cl\_is}) \quad [9]$$

and $EC50_{Screen/Loss}$ is right-shifted depending on the $Ca^{2+}$ levels present in solution, thus spanning the stability of the protein in the presence of $Ca^{2+}$:

$$EC50_{Loss} = EC50_{Loss\text{-no bivalents}} \cdot (1 + bivalent\_related\_folding) \quad [10]$$

Note that, depending on the levels of environmental $Ca^{2+}$, a right-shift occurs in *Screen/Loss* that leads to higher *FOF* as compared to environments where monovalent cations are present.

#### RESULTANT OF MODEL PROCESSES 1-4: CASQ2 fraction that is optimally folded

The fraction of CASQ2 that is optimally folded/functional (*FOF*) critically depends on ionic conditions, and is calculated by the equation:

$FOF = Promoted\_folding \cdot Loss$ [11]

Since the process of charge neutralization defines the EC50s for *Promoted_folding* and *Loss*, this process is fully present in equation [11] even if not mentioned explicitly.

### EQUILIBRIUM CONSTANTS OF THE CHEMICAL REACTION [1]:

*K1* ($D_i \rightleftharpoons 2M_i$) was defined to ensure that weak ionic strengths promote the formation of $D_{nf}$ whereas high ionic strengths (by rising the value of *K1*) lead to formation of $M_{nf}$ monomers.

$K1 = 1 / (1 + 0.25 \cdot \text{ionic\_strength})$ [12]

In order to assign a numeric value to *K2* ($2M_{nf} \rightleftharpoons 2M_o$), we made the conjecture that the folding-unfolding reaction of monomers is much faster than the formation of dimers, and that we start reaction [1] with all CASQ2 molecules being monomers. From the definition of $K2 = M_o / M_{nf}$ and these special conditions, we get that:

$K2 = FOF / (1 - FOF)$ [13]

*K3* ($2M_o \rightleftharpoons D_o$) was given a fixed numerical value. This was done because the experiments within the current manuscript demonstrate that CASQ2 dimers do occur independently of the presence of $Ca^{2+}$, and because no obvious dependency on KCl levels could be extracted from the data for this type of dimerization.

$K3 = 0.25$ [14]

$K_{tetramerization}$ was parametrized as being directly proportional to *bivalent_related_folding* (i.e. the $Ca^{2+}$ contribution to cationic ionic strength; see [9]) and inversely proportional to the $4^{th}$ power of *Loss* (see [8]). This definition is purely phenomenological, reflecting the facts that (a) CASQ2 does not form multimers in the absence of divalent cations (Supplementary Figure 6 in the manuscript) and that (b) polymers are destabilized by ionic strengths above those needed for ζ-potential close to neutrality (descending curve in Figure 6A).

$K_{tetramerization} = bivalent\_related\_folding \cdot Loss^{\,4}$ [15]

Finally, α was given a fixed numerical value.

$\alpha = 1.1$ [16]

Note that, despite the numerical value of α is small, the value of the equilibrium constants for polymerization reactions follow an exponential dependency on the number of dimers involved (see [3]). Therefore, such equilibrium constants can reach large values when the number of dimers is large. For example, when $n = 20$ dimers/polymer, the equilibrium constant becomes five times and a half the tetramerization constant

$K_{n=20} = K_{tetramerization} \cdot \alpha^{18} = K_{tetramerization} \cdot 5.56$

### SIMULATION METHODOLOGY

All simulations were performed in MATLAB R2021b (Mathworks Inc., Natick, MA, United States) on a Windows 11 Pro HD desktop computer. The function *fsolve* was called to solve for $[M_o]$, with options 'TolFun' =1e-12 and 'TolX' = 1e-12. The initial guess used by *fsolve* was $[M_o]_{initial\_guess} = [CASQ2_{Total}] \cdot FOF$. Calsequestrin 2 total concentration is input in the model in μM units, while ionic concentrations are input in mM units. The value of the equilibrium constants varies with each environment tested, according to the equations and formulas discussed above.

Each specific environment can be tested within a few seconds, so that complete sets of simulations can be performed in less than a minute.

Use of artificial intelligence: Meta AI (Meta, USA) was used to device the MATLAB code that calculates the molecular weight and the net charge (pH 7.3) of the CASQ2 monomer.

### MODEL OUTCOMES

#### Model Outcome 1: Ionic environments needed for zero ζ-Potential

When the model replicates the ionic environments used in the experiments, the application of equations [4] and [5] (setting "Non-neutralized charges = 0 %") renders that the ionic concentrations needed to neutralize CASQ2 approximate the observed ionic conditions experimentally needed to obtain a ζ-potential ≈ 0 (Table 1).

| Composition of ionic environment | MS. Fig. | Observed: [Salt] for ζ-potential ≈ 0 | Predicted by model: [Salt] needed to neutralize CASQ2 |
|---|---|---|---|
| Variable (1 to 600 mM) KCl | 2D | 194.2 mM KCl | Same as observed, since this value was used to obtain X in eq. [5] |
| Variable (1 to 600 mM) NaCl | | 159.7 mM NaCl | 139 mM NaCl |
| 50 mM KCl, variable (1 to 40 mM) $CaCl_2$ | 5C | 35.2 mM $CaCl_2$ | 35.3 mM $CaCl_2$ |
| 150 mM KCl, variable (1 to 40 mM) $CaCl_2$ | | 10 mM $CaCl_2$ | 10.8 mM $CaCl_2$ |

**Table 1: Observed vs. calculated salt concentrations at which z-potential ≈ 0 in different ionic environments.**

From the table it is clear that the theory is in excellent agreement with the experiment on what pertains the CASQ2 behavior in environments containing $K^+$ and $Ca^{2+}$. In environments containing solely NaCl, the theory some refinements may be necessary to fully account for the CASQ2 behavior.

#### Model Outcome 2: Apparent molecular weight of CASQ2 monomers as a function of [KCl]

The molecular weight (MW) of individual amino-acids was obtained from [1,2] and added, using the canonical human CASQ2 sequence (Uniprot ID O14958-1). Using this approach, it was obtained that the MW of CASQ2 monomers is 46.407 KDa. The weights used for each amino-acid were, in summarized form:

Amino-acid weights = {'A', 'R', 'N', 'D', 'C', 'E', 'Q', 'G', 'H', 'I', 'L', 'K', 'M', 'F', 'P', 'S', 'T', 'W', 'Y', 'V'}, [71.03711, 156.10111, 114.04293, 115.02694, 103.00919, 129.04259, 128.05858, 57.02146, 137.05891, 113.08406, 113.08406, 128.09496, 131.04049, 147.06841, 97.05276, 87.03203, 101.04768, 186.07931, 163.06333, 99.06841]

To explain the experimentally observed changes in the CASQ2's apparent molecular weight, it was assumed that optimal folding of the CASQ2 monomer leads to a reduction of its apparent molecular weight, following a simple linear relationship.

$$app\_MW = MW - \text{apparent\_compaction\_due\_to\_proper\_folding} * FOF \qquad [16]$$

where *app_MW* is the apparent molecular weight of the CASQ2 monomer, *MW* is the molecular weight of the unfolded CASQ2 monomer as calculated from its constituent amino-acids (see above), and apparent_compaction_due_to_proper_folding = 13 KDa.

Figure A.1, panel A, displays the processes used by the model to calculate *FOF* (equations [5] to [11]), whereas panel B compares the output of the model to Figure 2A within the manuscript. The qualitative agreement with the experimental data is remarkable.

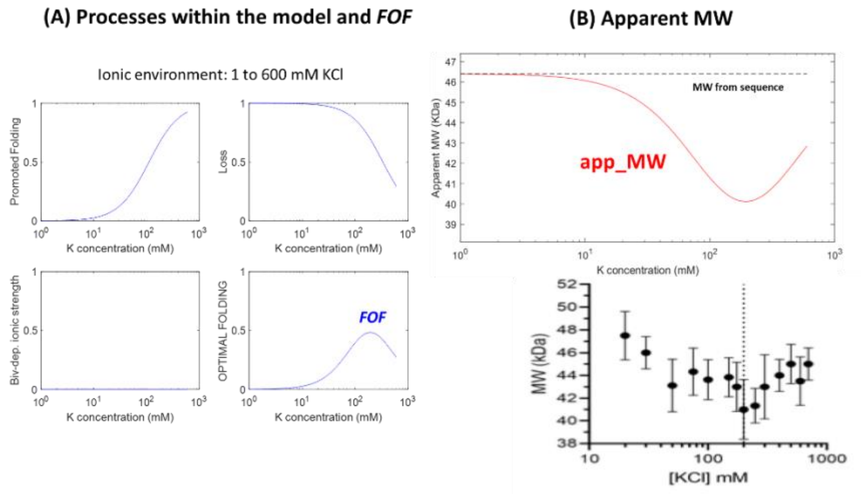


**Figure A.1: Apparent molecular weight of CASQ2 monomers in variable [KCl].**

**Model Outcome 3: Apparent molecular weight of CASQ2 monomers as a function of [$CaCl_2$] at different ionic strengths**

Figure S2 within the manuscript shows that, when CASQ2 dwells in an environment with 50 mM KCl, the addition of 1 to 10 mM $CaCl_2$ leads to an increase in ionic strength which itself leads to molecular compaction of the CASQ2 monomer. This effect is not observed in 200 mM KCl, where molecular compaction is already maximal.

In order to understand whether the current theoretical formulation can also simulate this behavior, we performed simulations as in *Model outcome 1* above, adjusting the concentrations of KCl and $CaCl_2$ to match the different experimental environments. Results are summarized by Figure A.2: Panel A.2-left replicates Figure S2 from the manuscript, whereas panels A.2-center and A.2-right compare, respectively, the theoretical predictions in 50 mM vs. 200 mM KCl. It can be seen that, as occurs in the experiments, a monomer compaction progressively develops with increasing levels of $Ca^{2+}$ in 50 mM KCl, whereas at 200 KCl the compaction of the CASQ2 monomer is already completed.

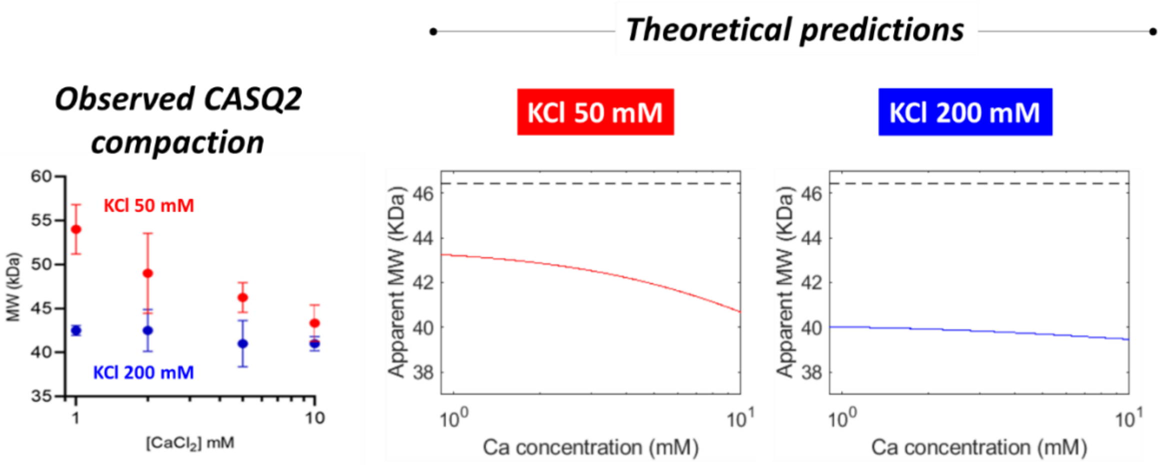


**Figure A.2: Apparent molecular weight of CASQ2 monomers in variable [$CaCl_2$] at low vs. optimal ionic strengths.**

**Model Output 4: [Dimer]-to-[Monomer] (D/M) ratio as a function of [KCl]**

Figure 3A within the manuscript shows that a non-negligible proportion of CASQ2 dimers are observed at any [KCl] in the absence of bivalents at [$CASQ2_{Total}$] = 2.5 μM. Furthermore, no polymers were detected using turbidometry (Figure S6). The D/M ratio is highest when the ionic strength is lowest, suggesting that the excess of dimers at low ionic strengths originates via non-specific electrostatic interactions due to incomplete charge shielding of the CASQ2 molecule.

To understand whether the current theory could simulate this behavior, $N$ = 20 was set in the model and [$CASQ2_{Total}$] = 2.5 μM. The processes and *FOF* calculated by the model are shown in Figure A.1A above. Figure A.3A shows the concentration of the different chemical species, whereas Figure A.3B compares the predicted D/M ratio vs. Figure 3A within the manuscript. The qualitative agreement between theory and experiment is close, and adds value to the hypotheses presented in the manuscript: (a) at low ionic strengths and in the absence of bivalents, most dimers are non-functionally folded; (b) at high ionic strengths and in the absence of bivalents, the dimers observed are mostly folded in an optimal (i.e. polymerization-ready) fashion. No polymers are observed.

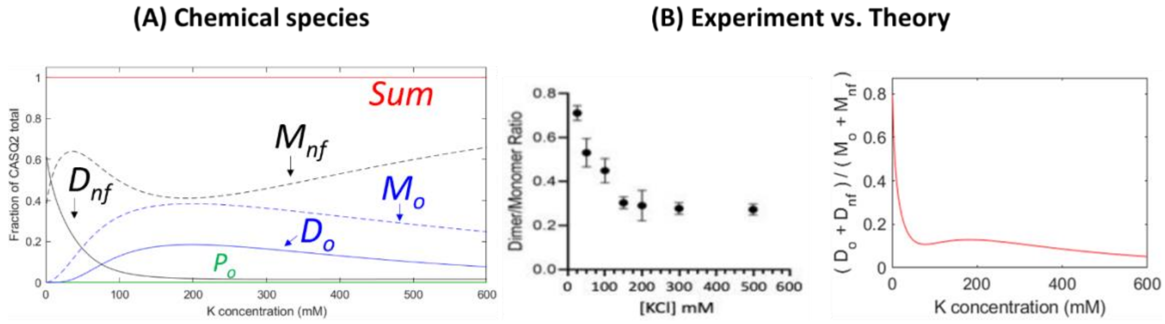


**Figure A.3: Behavior of the [Dimer]-to-[Monomer] in variable [KCl] and no added $Ca^{2+}$.**

**Model Output 5: [Dimer]-to-[Monomer] ratio as a function of [$CaCl_2$] in environments with 50 mM [KCl]**

Figure 5A within the manuscript shows that, at [KCl] = 50 mM, the addition of $CaCl_2$ (1-60 mM) leads to non-monotonic behaviors of the D/M ratio. We tested this behavior by setting in the model [KCl] = 50 mM, $N$ = 20 and [$CASQ2_{Total}$] = 2.5 μM.

Figure A.4A shows the processes within the model, Figure A.4B shows the chemical species and Figure A.4C compares the experimental observation in the manuscript (Figure 4A) to the theoretical predictions. While the curve calculated by the model clearly fails in replicating the quantitative CASQ2 behavior, it replicates with excellent accuracy its qualitative behavior (expressed by the $Ca^{2+}$ levels at which the different experimental effects are observed): (a) 1-5 mM $Ca^{2+}$ rapidly leads to lesser fraction of non-functionally folded molecules in favor of optimally folded ones, decreasing the overall number of $D_{nf}$ dimers and the D/M ratio; (b) between 5 and 10 mM, the increase in optimally folded molecules is not yet high enough, and so the D/M ratio remains at a valley; (c) above 10 mM, the monomers and dimers that now predominate are optimally folded, and the D/M ratio rises above its value in 1 mM $Ca^{2+}$.

At 50 and 60 mM $Ca^{2+}$, the qualitative theoretical behavior deviates from the experiment, since in the theory the D/M ratio flattens. This fact, and the quantitative disagreement highlighted above, indicate that further refinements of the theory are needed when describing the CASQ2 behavior in the presence of bivalents. That said, it should also be noted that, at high $Ca^{2+}$ concentrations (50–60 mM), Mass Photometry measurements become intrinsically less reliable due to the concurrent formation of higher-order oligomers. This phenomenon, which also occurs at elevated $K^+$ concentrations, increases background noise and reduces the precision of assignment of counts to the dimeric species. Therefore, the apparent flattening of the experimental D/M ratio at high $Ca^{2+}$ levels may also reflect, at least in part, methodological limitations rather than a genuine deviation from the theoretical prediction.

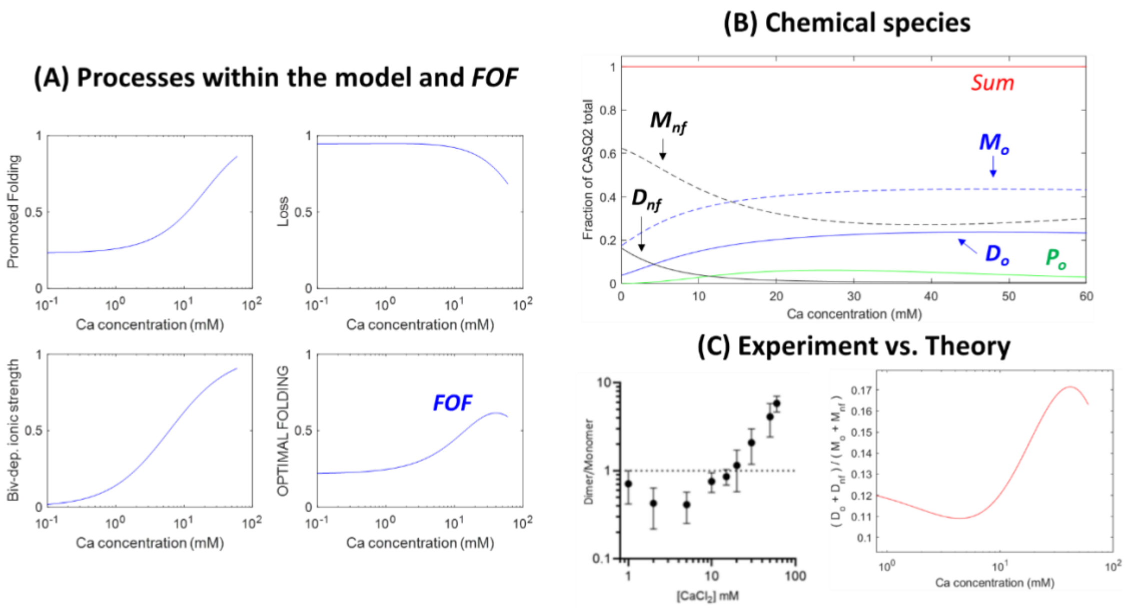


**Figure A.4: Behavior of the [Dimer]-to-[Monomer] in 50 mM [KCl].**

**Model Output 6: [Dimer]-to-[Monomer] ratio as a function of [$CaCl_2$] in 50 vs. 150 mM [KCl]**

Figure 4B within the manuscript shows that, when KCl concentration equals 150 mM, the $Ca^{2+}$-dependent decrease in D/M ratio (seen in 50 mM KCl at low millimolar $Ca^{2+}$) flattens due to attenuation of the $K^+$ vs. $Ca^{2+}$competition in favor of potassium. To see whether the theory could qualitatively replicate this behavior, we performed new simulations by setting in the model [KCl] = 150 mM, N = 20 and [$CASQ2_{Total}$] = 2.5 μM

Figure A.5 compares the theoretical D/M ratios at 50 mM (left) and 150 mM KCl (center) vs the experiments shown in Figure 4B of the manuscript (right). The model qualitatively replicates the behavioral shift shown by the experiments (note arrowheads inserted within the theoretical predictions).

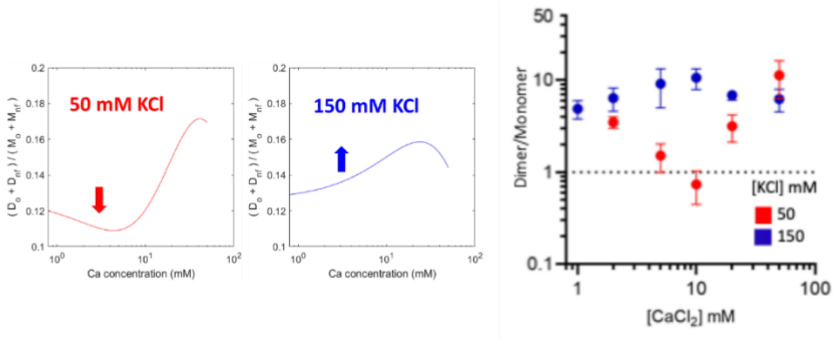


**Figure A.5: Behavior of the [Dimer]-to-[Monomer] ratio in 50 mM vs. 150 mM [KCl]**

**Model Output 7: CASQ2 polymerization is a $Ca^{2+}$-dependent switch in which $K^+$ shows mixed effects**

The experiments shown within the current manuscript are aimed at studying the $K^+$ vs. $Ca^{2+}$ relationship in shaping CASQ2 polymerization. 3 effects are clearly seen:

- In the absence of $Ca^{2+}$ ions, $K^+$ ions alone are not capable of leading to CASQ2 polymerization, regardless of their amounts (Figure S6)
- CASQ2 polymerization starts at lower $Ca^{2+}$ concentrations when $K^+$ levels are higher (a sensitizing effect on the part of $K^+$) (Figure 6A)
- At high $K^+$ levels, $Ca^{2+}$-dependent CASQ2 polymerization displays a lower cooperativity (a negative effect on the part of $K^+$) (Figure 6A)

The lack of polymerization in the absence of $Ca^{2+}$ can be already been seen in the simulations from Figure A.2 ($[CASQ2_{Total}]$ = 2.5 µM).

To simulate the two latter CASQ2 behaviors, we set $[CASQ2_{Total}]$ = 45 µM (as in the experiments) and $N = 20$. Two KCl-containing environments were simulated (either 10 mM or 50 mM, as in the experiment), to which $Ca^{2+}$ ions were added within the concentration range 1-200 mM. Figure A.5, left, replicates the experiments shown in Figure 6A of the manuscript. Theoretical results are shown to the right. The green traces in the right (theoretical) panels, which indicate the compounded sum of tetramers, hexamers, octamers, etc (up to 40 monomers) represent well the experimental behavior: there are both an ascending and a descending phase. The model replicates both the lower threshold $[Ca^{2+}]$ and the smallest steepness of the polymerization curve in 50 mM vs. 10 mM KCl. Of additional importance is the formation of $M_{nf}$ monomers and $D_o$ dimers concomitantly to the dissipation of polymers during the descending phase of the curve, implying that polymer folding has become impeded by the high ionic strength. The model, however, right-shifts the descending phase of both polymerization curves relative to those of the experiment, and it right-shifts the ascending curve of the 10 mM KCl environment, indicating once again that theory refinement is needed.

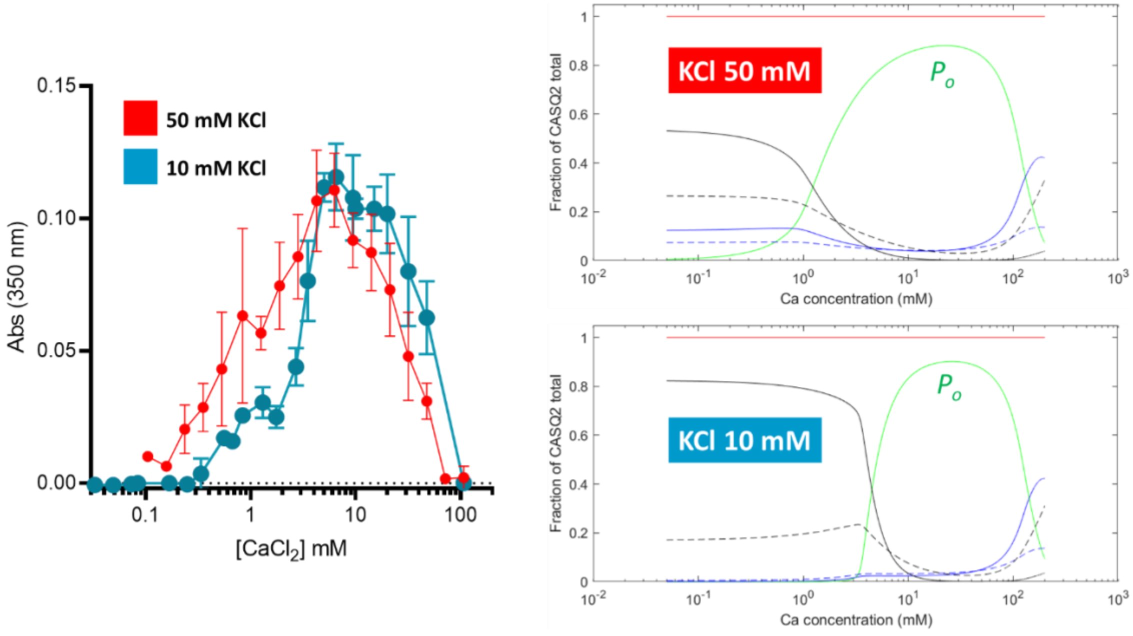


**Figure A.6: $Ca^{2+}$-dependent CASQ2 polymerization (green line) in environments with low [KCl].**

**Model Output 8: $Ca^{2+}$-dependent CASQ2 polymerization occurs suddenly**

Figure 7A within the manuscript highlights the $Ca^{2+}$-dependent switch-like behavior of CASQ2 polymerization, by testing the size of CASQ2 particles at either 50 or 150 mM environmental KCl. It is observed that polymerization occurs suddenly, indicating a switch-like event of extreme cooperativity, which threshold [$Ca^{2+}$] depends on [$K^+$]. The experiments were not conducted beyond 40 mM $Ca^{2+}$, so a descending phase cannot be visualized.

We tested whether the theory can simulate this behavior (Figure A.6) by setting [$CASQ2_{Total}$] = 45 µM and $N$ = 20. The left panel in Figure A.6 is the experiment. The central and right panels are the predicted CASQ2 behaviors at, respectively, 50 and 150 mM KCl. Within these theoretical behavior panels, the graphs separately display the individual fractions of $M_o$, $D_o$ and each species $P_o$, up to a [$Ca^{2+}$] = 15 mM, for any given environmental [$Ca^{2+}$]. It can be seen that, in both environments, a polymerization reaction occurs suddenly, with few polymer species in-between the size of the dimer (2 $M_o$/polymer) and the maximum size allowed (40 $M_o$/polymer). Furthermore, for the 50 mM KCl environment the formation of very large polymers is about 75% ended at 5 mM $Ca^{2+}$, closely resembling the experiment. For the 150 mM KCl environment, formation of very large polymers starts around 3-4 mM $Ca^{2+}$, and by a level of 10 mM such polymerization is nearly completed. Again, no polymers of intermediate sizes are observed.

It can be concluded that the theoretical 0-D CASQ2 behavior, as simulated herein, reproduces the experimental observation by which a $Ca^{2+}$-dependent switch triggers the formation of very large polymers in a cooperative fashion.

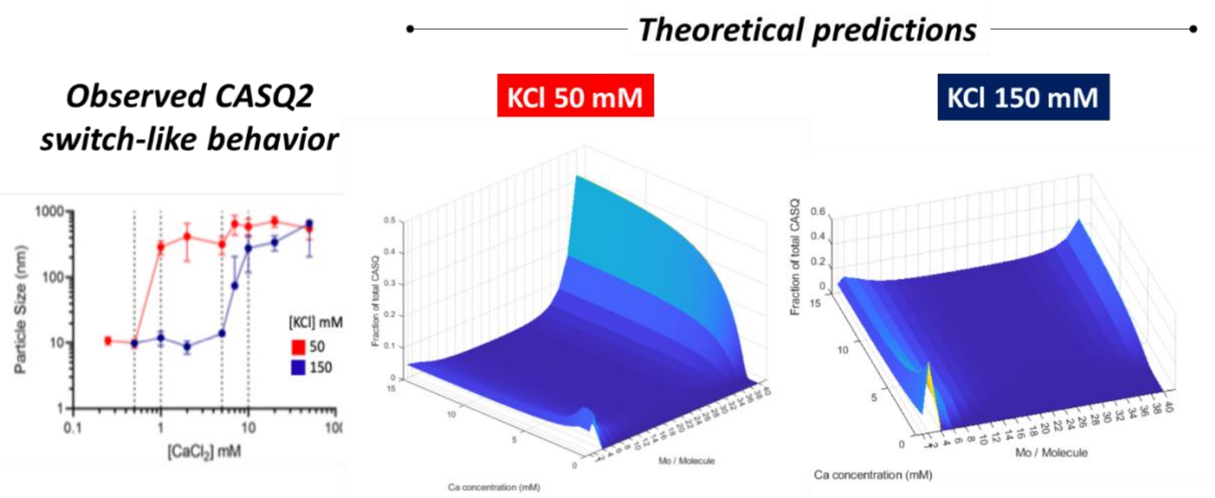


**Figure A.7: CASQ2 polymerization occurs suddenly**

### CONSISTENCY OF THE 0-D CASQ2 MODEL vs. PREVIOUSLY PUBLISHED DATA

The 0-D theoretical formulation presented herein focuses solely in the equilibrium folding behavior of CASQ2. This leaves unanswered the key question of what happens to $Ca^{2+}$ binding by CASQ2, and in particular the question of whether the current theoretical formulation is (or not) consistent with previously published $Ca^{2+}$ binding experiments. Note that the current manuscript (and this appendix) present evidence suggesting that CASQ2 is able to dimerize even in the absence of $Ca^{2+}$. This is, however, apparently contradictory to experiments such as those from Park et al. (2004) [4], which were interpreted to indicate that there is a strict $M_o \rightarrow D_o \rightarrow P_o$ sequence, and that dimerization is very strongly dependent on [$Ca^{2+}$]. Therefore, the question arises: Is the current theory capable of reconciling the experiments from both publications and provide a unified interpretation?

In order to address the question, it was first assumed that $Ca^{2+}$ binding to CASQ2 can only occur in optimally folded molecules (i.e. $M_o$, $D_o$ and $P_o$ of any size). Maximum $Ca^{2+}$-binding capacity of CASQ2 was set to 47.45, obtained as explained in Footnote 1. It was assumed that $Ca^{2+}$-binding capacity was independent of polymer size, since the theory has no way of calculating bound $Ca^{2+}$ other than based on the charge neutralization principle (see Footnote 1). Therefore, the role of $Ca^{2+}$ as a bridging element between monomers has been omitted in this calculation. All the above implies that:

binding_sites_monomer = 47.45

binding_sites_dimer = 2 · n_binding_sites_monomer

binding_sites_polymer$_n$ = $n$ dimers · binding_sites_dimer

where $n$ is the number of dimers in the polymer of size $n$. Dissociations constants were set, for the monomer, $K_{D\text{-}mono}$ = 0.25 mM, and for species with 2 or more monomers, $K_{D\text{-}higher}$ = 5 mM. Hill coefficients were assumed to be $H_{mono}$ = 1 and $H_{high}$ = 4, respectively, for monomers or species with more than one monomer. Occupancy was defined as the moles of $Ca^{2+}$ bound per total CASQ2 present, as in Park et al. In the simulations, $[CASQ2_{Total}]$ was 45 µM (about 2 mg/ml, mid-range of Park et al. experiments) and $N$, the maximum number of dimers allowed within individual polymers, was 20. The environment was set to [KCl] = 300 mM, as in Park et al. We did not modify the pH of our simulation (7.3), which is slightly different from that of Park et al. (7.5).

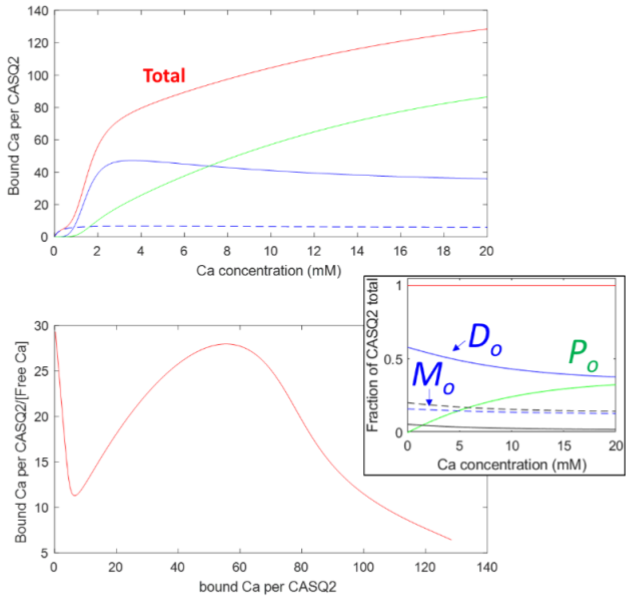


**Figure A.8: $Ca^{2+}$-binding properties of 0-D simulated CASQ2 as a function of free $[Ca^{2+}]$**

Figure A.8 shows the theoretical prediction. The top and bottom panels were built similarly to Figures 5A (binding curve) and 5B (Scatchard-like plot) from Park et al. (2004) [4]. An insert further contains the proportion of $D_{nf}$, $M_{nf}$, $M_o$, $D_o$, and all combined $P_o$ molecules (note that the proportion of $D_o$ dimers, as predicted by the current 0-D electrostatic theory, is very large even in the absence of $Ca^{2+}$). Within the top panel of Figure A.8, the dashed blue line represents $Ca^{2+}$ that is bound to CASQ2 $M_o$ monomers, whereas the solid blue line is $Ca^{2+}$ bound to $D_o$ dimers. The green line is the combined $Ca^{2+}$ that is bound to all other polymers (i.e. tetramers, hexamers, octamers, and so on). Finally, the red curve labelled with "Total" is the sum of the above 3 curves, and the one that needs to be compared to Figure 5 in Park et al. (2004).

There is a good qualitative agreement: It is evident in the binding curve that sequential binding phases occur (first to monomers, then to dimers, then to polymers). Furthermore, the theoretical Scatchard-like plot shows a similar behavior vs. the one observed experimentally, with varying slopes as a function of occupancy. However, what explains the different phases according to the current theory are key differences in $K_D$, Hill coefficient and monomer/dimer/polymer abundances, rather than a strict coupling between binding and polymerization.

Areas of disagreement include the maximum $Ca^{2+}$ bound ions per CASQ2 molecule, with was shown experimentally to be 36 by Park et al., vs. 47.45 ions calculated by the 0-D theory. This is in part because the 0-D theory only assumes electrostatics, without structural distinctions of what constitutes (or not) a $Ca^{2+}$-binding site in three-dimensional space.

All the above said, we conclude that our theoretical frame can qualitatively replicate both the experiments from the current manuscript and those from Park et al. (2004). Therefore, we see no contradiction among the two sets of experimental data.

### THE NEAR-PHYSIOLOGICAL ENVIRONMENT

As a last simulation, we applied the current theory to understand what would be the expected aggregation status of CASQ2 within environments as close as possible to the physiological cardiac environment. We set as near-physiological conditions 140 mM KCl, 5 mM NaCl and variable free $Ca^{2+}$, ranging from 0 to 1.25 mM. [$CASQ2_{Total}$] was set to 1000 μM, the concentration believed to occur within the junctional sarcoplasmic reticulum of striated muscle [5]. $N$ was set to 20, as in prior simulations.

Figure A.9 summarizes the theoretical expectation. The left panel indicates that at free $Ca^{2+}$ levels of 1 mM (i.e. similar to levels found in the junctional sarcoplasmic reticulum during diastole) approximately 5% of the total CASQ2 would be in the form of $D_o$ dimers, whereas most CASQ2 (about 95%) would be expected to form oligomers and high-order polymers. A very small fraction (less than 1%) would be monomeric, of either optimal or non-functional configurations. Figure A.9, right panel, dissects which optimally folded forms would predominate: the vast majority of CASQ2 protein (about 90%) would form the highest polymers allowed by the simulation, whereas the remaining fraction would form tetramers, hexamers and octamers, with few polymers of intermediate order. Together, the theoretical predictions agree well with the observation (by electron microscopy) of large CASQ2 polymers in the junctional sarcoplasmic reticulum from wild-type cells [6]. In diseases such as Catecholaminergic Polymorphic Ventricular Tachycardia type 1, where the sarcoplasmic reticulum $Ca^{2+}$ content is similar or slightly reduced (due to $Ca^{2+}$ leak) vs. healthy cells, the theory still predicts that CASQ2 will be highly polymerized. This is in agreement with the visualization of polymers in electron micrographs from affected animals [7].

It would be tempting to speculate about the predictions at free $Ca^{2+}$ levels consistent with the nadir of sarcoplasmic reticulum depletions during a heartbeat. However, due to the equilibrium nature of the current theory, and because the time lapse where the sarcoplasmic reticulum remains depleted during a heartbeat is narrow, we prefer to refrain from such speculation.

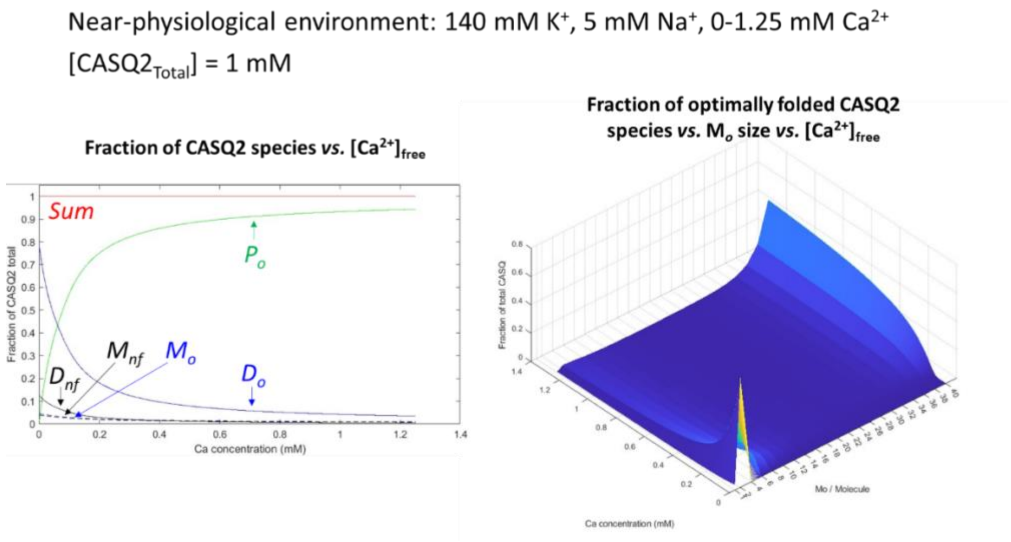


**Figure A.9: Predicted CASQ2 aggregation status in simulated near-physiological junctional sarcoplasmic reticulum ionic environment**

An additional question faced by the theory is, if the ionic environment within the entire sarcoplasmic reticulum is similar, why does CASQ2 not polymerize in the vicinity of the transcription sites [8]. We believe it is because, at transcription sites, the CASQ2 concentration is not high-enough to promote strong polymerization. We are aware of the fact that CASQ2 concentration may not be the only determinant of its polymerization state at transcription. However, this model is not currently suited for incorporating the effect of post-translational modifications. To illustrate this point, we have repeated the simulation shown by Figure A.9 but using an arbitrarily low [$CASQ2_{Total}$] value equal to 10 µM. Figure A.10, analogous in structure to Figure A.9, shows the results. Figure A.10, left, confirms the expectation by showing that at 1 mM free $Ca^{2+}$ the fraction of $P_o$ molecules is very small. Figure A.10, right, further shows that the totality of $P_o$ molecules are tetramers. While the levels of CASQ2 chosen for this simulation were arbitrarily set to illustrate a point, it is worth noting that the simulation outcome replicates the accumulation of tetrameric DsRed-CASQ2 at the CASQ2 transcription sites as shown by the Cala group [8].

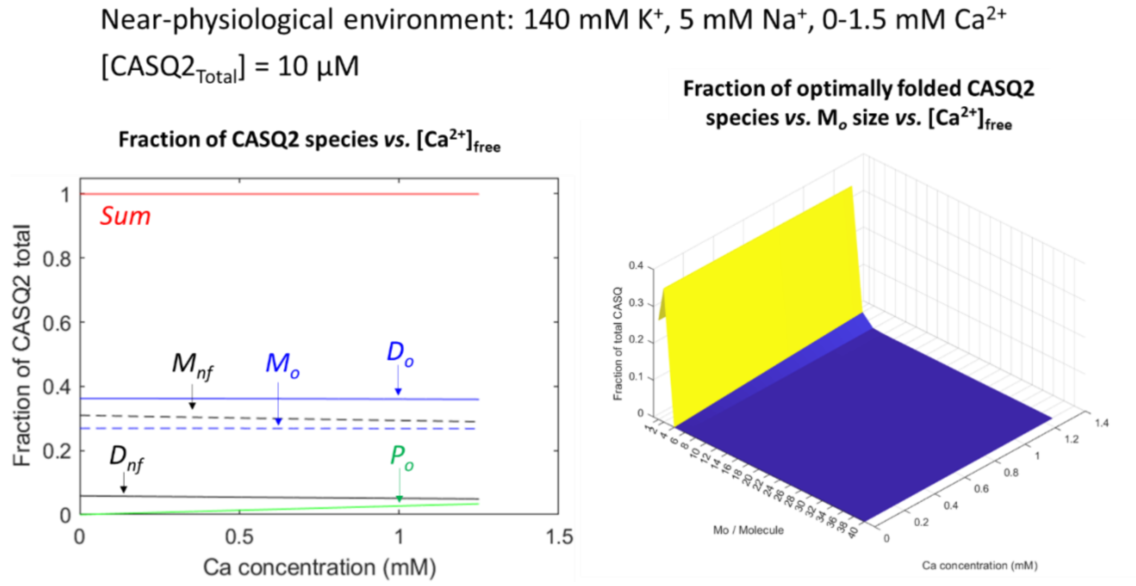

**Figure A.10: Predicted CASQ2 aggregation status in simulated near-physiological transcription site environment**

### LIMITATIONS

The 0-D nature of this theory allows for interested users to perform a very fast and efficient exploration of the CASQ2 behavior in a plethora of ionic environments, in such a way that it can be used predictively, at least in qualitative terms. However, this same 0-dimensionality creates difficulties when predicting the precise number of $Ca^{2+}$ binding sites, as it does not consider the 3-D structure of the protein: Bound $Ca^{2+}$ can only be calculated based on electrostatics (see Footnote 1), and, while the calculation agrees well with some experimental estimates [3], it does not agree with others [4]. One further limitation is that the theory does not contemplate the effects of post-transcriptional modifications, nor of $Mg^{2+}$-rich environments, on CASQ2 aggregation dynamics, since the experimental data that were used to create the theory were not generated in those conditions.

### CONCLUDING REMARKS

Despite being phenomenological and 0-dimensional, the theoretical formulation presented in this appendix retains a glimpse of physical intuition, is quite simple, and approximates very important qualitative *in vitro* behaviors of CASQ2, including monomer compaction, dimerization and polymerization behavior in different ionic environments, and $Ca^{2+}$ binding studies. Thus, this theoretical framework has the potential, if properly refined and extended, to aid in the scientific understanding about CASQ2 and $Ca^{2+}$-dependent cardiac arrhythmias.
